# Ancestry inference and grouping from principal component analysis of genetic data

**DOI:** 10.1101/2020.10.06.328203

**Authors:** Florian Privé

## Abstract

Here we propose a simple, robust and effective method for global ancestry inference and grouping from Principal Component Analysis (PCA) of genetic data. The proposed approach is particularly useful for methods that need to be applied in homogeneous samples. First, we show that Euclidean distances in the PCA space are proportional to *F*_*ST*_ between populations. Then, we show how to use this PCA-based distance to infer ancestry in the UK Biobank and the POPRES datasets. We propose two solutions, either relying on projection of PCs to reference populations such as from the 1000 Genomes Project, or by directly using the internal data. Finally, we conclude that our method and the community would benefit from having an easy access to a reference dataset with an even better coverage of the worldwide genetic diversity than the 1000 Genomes Project.

## Introduction

Principal Component Analysis (PCA) has been widely used to correct for population structure in association studies and has been shown to mirror geography in Europe (Price *et al.* 2006; Novembre *et al.* 2008). Due to its popularity, many methods has been developed for efficiently performing PCA (Abraham *et al.* 2017; Privé *et al.* 2020) as well as appropriately projecting samples onto a reference PCA space (Zhang *et al.* 2020; Privé *et al.* 2020), making possible to perform these analyses for ever increasing datasets in human genetics. Naturally, PCA has also been used for ancestry inference. However, among all studies where we have seen PCA used for ancestry inference, we have found there was no consensus on what is the most appropriate method for inferring ancestry using PCA. For example, there may be divergences on which distance metric to use and the number of PCs to use to compute these distances.

Here, we first compare several distance metrics with the popular *F*_*ST*_ statistic between populations. We show that the simple Euclidean distance on PC scores is the most appropriate distance to use, then we show how to use it to infer global ancestry and to group individuals in homogeneous sub-populations. We do not provide a method to infer admixture coefficients nor local ancestry, which are different problems for which there are several existing methods (Alexander *et al.* 2009; Frichot *et al.* 2014; Raj *et al.* 2014; Padhukasahasram 2014). However, inferring global ancestry in non-admixed individuals is still a very important problem since there are methods that specifically need to be applied in samples of homogeneous ancestry. This is the case e.g. for polygenic score methods that have been shown to underperform when applied to populations not homogeneous to the population used for training (Martin *et al.* 2017).

## Measures of genetic dissimilarity between populations

We first compare four measures of genetic dissimilarity using populations of the 1000 Genomes Project (1000G^†^, 1000 Genomes Project Consortium *et al.* (2015)). The *F*_*ST*_^†^ is an ubiquitous measure of genetic dissimilarity between populations and the first measure we use in this comparison. We report *F*_*ST*_ between the 26 1000G populations in tables S1-S5, and the clustering of these populations based on *F*_*ST*_ in figure S1. The other three measures compared are distances applied to the PC scores^†^ of the genetic data: 1) the Bhattacharyya distance^†^; 2) the distance between the centers (geometric medians^†^) of the two populations; 3) the shortest distance between pairs of PC scores from the two populations. The (squared) Euclidean distance between population centers appears to be an appropriate PCA-based distance as it is approximately proportional to the *F*_*ST*_ (Figure 1) and provides an appropriate clustering of populations (Figure S4). In contrast, the two other Bhattacharyya and shortest distances do not provide as satisfactory results (Figures S2, S3, S6 and S7). For example, African Caribbeans in Barbados (ACB) and Americans of African Ancestry in SW USA (ASW) and the four admixed American (AMR) populations are close to all European (EUR), South Asian (SAS) and African (AFR) populations when using the Bhattacharyya distance (Figure S2). Using the shortest distances between pairs of individuals in two different populations is very sensitive to outliers. We also vary the number of PCs used for computing the Euclidean distances and how they compare with *F*_*ST*_ in figure S5. With 2 to 4 PCs, we are able to adequately separate distant populations, but not the closest ones. For example, when using 4 PCs, there are pairs of populations with an *F*_*ST*_ of ~0.02 while their PC centers are superimposed (Figure S5). When using more PCs (8, 16 or 25) to compute the distances, results remain similar.

**Figure 1:**
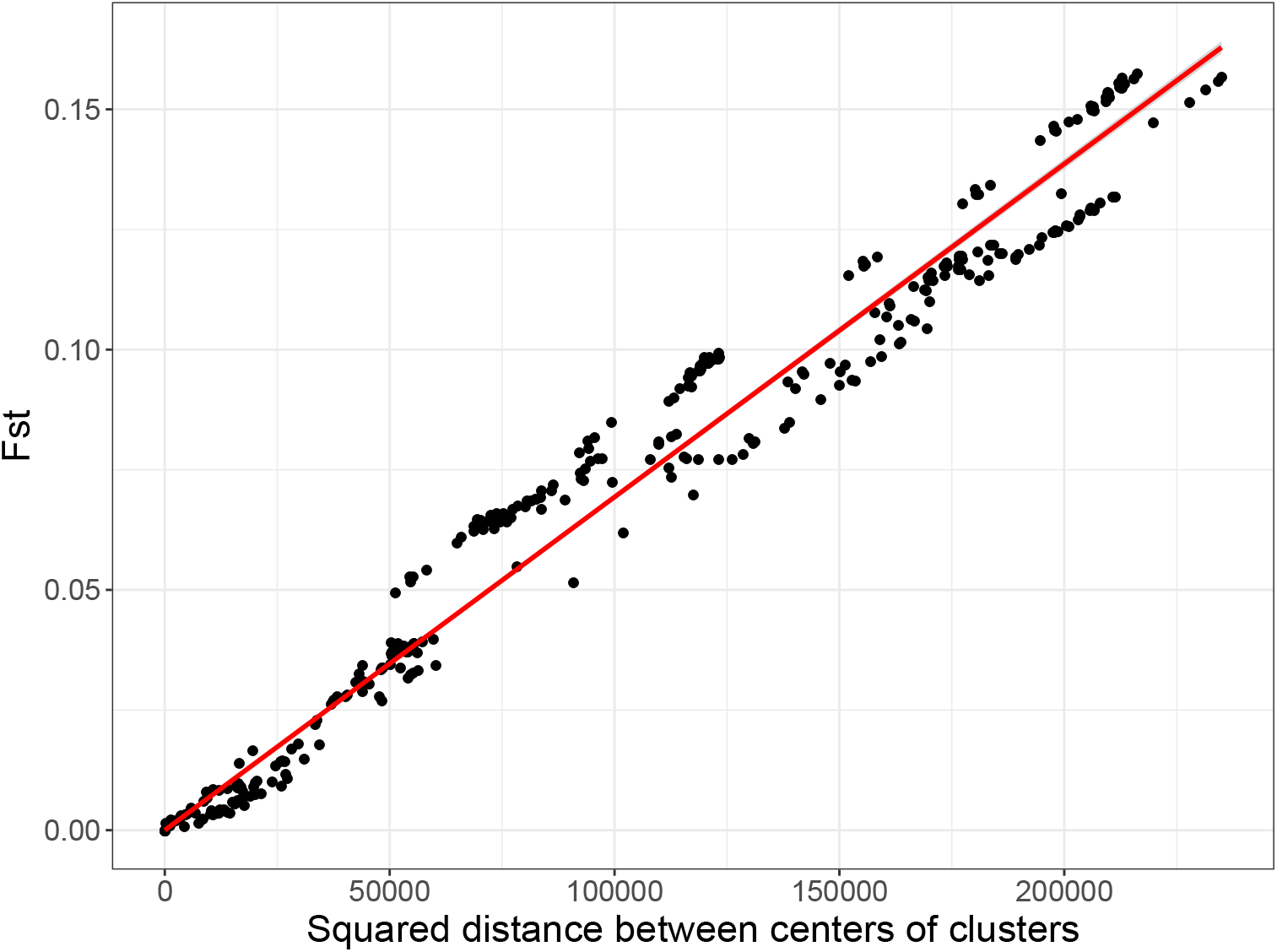
Comparing *F*_*ST*_ to the squared Euclidean distance on the PCA space between centers of pairs of the 26 1000G populations.

## PCA-based ancestry inference

We project the dataset of interest onto the PCA space of the 1000G data using the fast tools developed in Privé *et al.* (2020). We recall that this uses an automatic removal of LD when computing PCA and a correction for shrinkage in projected PC scores, which has been shown to be particularly important when using PC scores for ancestry estimation (Zhang *et al.* 2020). Based on the results from the previous section, we propose to assign individual ancestry to one of the 26 1000G populations based on the Euclidean distance to these reference population centers (geometric medians^†^) in the PCA space. Since we showed previously that (squared) distances in the PCA space are proportional to *F*_*ST*_, we can set a threshold on these distances that would correspond approximately to an *F*_*ST*_ of e.g. 0.002. This threshold is close to the dissimilarity between Spanish and Italian people (*F*_*ST*_ (IBS, TSI) of 0.0015). When an individual is not close enough to any of the 26 1000G populations, we leave its ancestry inference as unknown, otherwise we assign this individual to the closest reference population center.

We first perform ancestry estimation for the individuals in the UK Biobank^†^. These individuals seem to originate from many parts of the world when we project onto the PCA space of the 1000G (Figure S8). Self-reported ancestry (Field 21000) is available for almost all individuals, with only 1.6% with unknown or mixed ancestry. When using the threshold defined before, we could not infer ancestry for 4.6% of all 488,371 individuals. More precisely, among “British”, “Irish” and “White” ancestries, this represented respectively 2.2%, 3.3% and 7.9% (Tables 1 and S7). This also represented 3.3% for “Chinese”, 13.8% for “Indian” and 17.8% for “African” ancestries. Finally, mixed ancestries were particularly difficult to match to any of the 1000G populations, e.g. 97.3% unmatched within “White and Black Africa” and 93.0% within “White and Asian” ancestries. Only 47 individuals were misclassified in “super” population of the 1000G; e.g. six “British” were classified as South Asians, one “Chinese” as European and 25 “Caribbean” as South Asian by our method (Table 1). However, when comparing the location of these mismatched individuals to the rest of individuals on the PCA space computed within the UK Biobank (Bycroft *et al.* 2018), it seems more probable that our genetic ancestry estimate is exact while the self-reported ancestry is not matching the underlying genetic ancestry for these individuals (Figure S9). This possible discrepancy between self-reported ancestry and genetic ancestry has been reported before (Mersha and Abebe 2015).

**Table 1:** Self-reported ancestry (left) of UKBB individuals and their matching to 1000G continental populations (top) by our method. See the description of 1000G populations at https://www.internationalgenome.org/category/population/.

|  | AFR | AMR | EAS | EUR | SAS | Not matched |
| --- | --- | --- | --- | --- | --- | --- |
| British | 2 |  | 1 | 421457 | 6 | 9548 |
| Irish |  |  |  | 12328 |  | 425 |
| White | 1 | 1 | 1 | 499 |  | 43 |
| Other White |  | 40 |  | 11334 | 1 | 4440 |
| Indian |  |  |  | 5 | 4922 | 789 |
| Pakistani |  |  |  |  | 1421 | 327 |
| Bangladeshi |  |  |  |  | 217 | 4 |
| Chinese |  |  | 1453 | 1 |  | 50 |
| Other Asian | 1 |  | 279 |  | 939 | 528 |
| Caribbean | 3848 |  |  |  | 25 | 424 |
| African | 2633 |  |  | 1 |  | 570 |
| Other Black | 74 |  |  |  | 2 | 42 |
| Asian or Asian British |  |  | 2 |  | 20 | 20 |
| Black or Black British | 20 |  |  | 2 |  | 4 |
| White and Black Caribbean | 24 | 1 |  | 8 | 1 | 563 |
| White and Black African | 5 |  |  | 6 |  | 391 |
| White and Asian |  | 1 | 2 | 27 | 26 | 746 |
| Unknown | 835 | 173 | 576 | 2296 | 633 | 3307 |

We also test the approach proposed in *Zhang et al.* (2020) which consists in finding the 20 nearest neighbors in 1000G and computing the frequency of (super) population membership, weighted by the inverse distance to these 20 closest 1000G individuals. When this probability is less than 0.875, they leave the ancestry as unknown, aiming at discarding admixed individuals. Less than 0.5% could not be matched by their method (Table S6). Of note, they could match much more admixed individuals, whereas they set a high probability threshold aiming at discarding such admixed individuals. Morever, there are many more discrepancies between their method and the self-reported ancestry in the UK Biobank (Table S6) compared to the previous results with our method (Table 1). Finally, our method is able to accurately differentiate between sub-continental populations such as differentiating between Pakistani, Bangladeshi and Chinese people (Table S7). We also applied our ancestry detection technique to the European individuals of the POPRES data (Nelson *et al.* 2008). Only 16 out of the 1385 individuals (1.2%) could not be matched, of which 11 were from East or South-East Europe (Table S8). Note that all individuals that we could match were identified as of European ancestry. We could also identify accurately sub-regions of Europe; e.g. 261 out of 264 Spanish and Portugese individuals were identified as “Iberian Population in Spain” (EUR_IBS, Table S8).

## PCA-based ancestry grouping

Finally, we show several ways how to use our ancestry inference method for grouping genetically homogeneous individuals. One first possible approach is to simply match individuals that are close enough to one of the 1000G populations, as described previously. Alternatively, one could use the internal PC scores and the self-reported ancestries or countries of birth, e.g. available in the UK Biobank (Fields 21000 and 20115). This solution does not require projecting individuals to the 1000G, but does require computing PC scores within the dataset instead. In the UK Biobank data, we can define centers of the seven self-reported ancestry groups: British, Indian, Pakistani, Bangladeshi, Chinese, Caribbean and African; then match all individuals to one of these centers (or none if an individual is far from all centers). This enables e.g. to capture a larger set of individuals who are close enough to British people (e.g. Irish people), while discarding individuals whose genetic ancestry is not matching the self-reported ancestry (Table S9). Only 3.7% of all individuals could not be matched. The resulting clusters are presented in the PCA space in figure 2.

**Figure 2:**
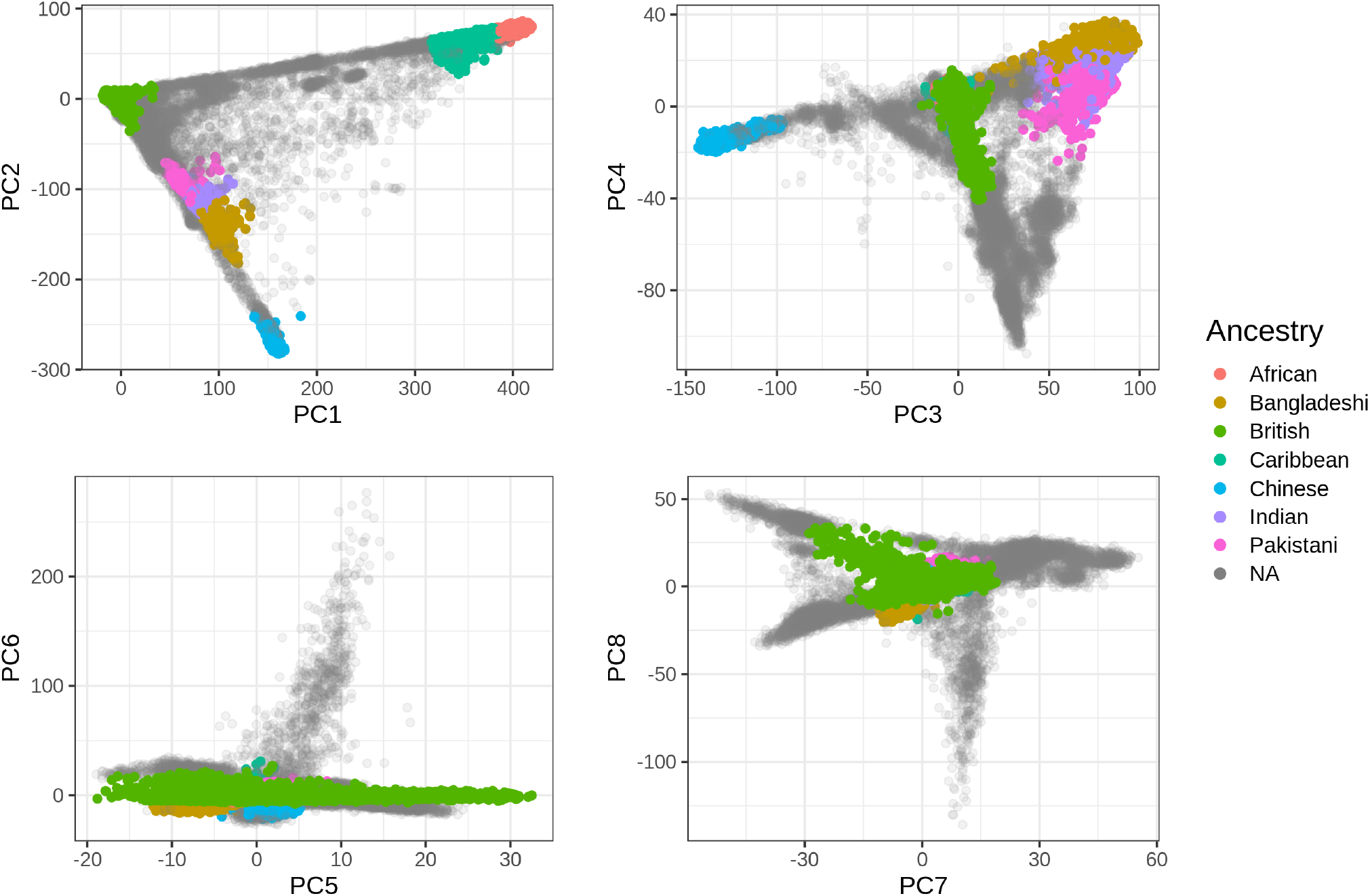
The first eight PC scores computed from the UK Biobank (Field 22009) colored by the homogeneous ancestry group we infer for these individuals.

One could do the same using the countries of birth instead of the self-reported ancestries. Again, the country of birth may sometime not reflect the ancestral origin. Therefore, we first compute the robust centers (geometric medians) of all countries with at least 300 individuals. Then, we cluster these countries based on their distance in the PCA space to make sure of their validity as proxies for genetic ancestry and to choose a small subset of centers with good coverage of the overall dissimilarities (Figure S10). Based on the previous clustering and the available sample sizes, we chosed to use the centers from the following eight countries as reference: the United Kingdom, Poland, Iran, Italy, India, China, “Caribbean” and Nigeria. Only 2.8% of all individuals could not be matched (Table S10). The resulting clusters are presented in the PCA space in figure S11. Note that these clusters probably include individuals from nearby countries as well.

Finally, when we know that the data is composed of a predominant ancestry, we can define a single homogeneous cluster by simply restricting to individuals who are close enough to the overall center of all individuals (Figure S12). When doing so, we can cluster 91% of the data into one cluster composed of 421,871 British, 12,039 Irish, 8351 “Other White”, 1814 individuals of unknown ancestry, 467 “White” and 41 individuals of other self-reported ancestries. This is made possible because we use the geometric median which is robust to outliers.

## Discussion

Here we propose a PCA-based method for ancestry inference and grouping individuals into genetically homogeneous clusters. We show how the PCA-based distance is related to the *F*_*ST*_, which allows to compute distances based on PC scores directly. This relation between *F*_*ST*_ and (squared) Euclidean distances in the PCA space has been previously shown for two populations only (McVean 2009). Previously, we and others proposed to use (robust) Mahalanobis distances to infer ancestry or identify a single homogeneous group of individuals (Peterson *et al.* 2017; Privé *et al.* 2020). When looking at distances between two populations, this corresponds to using the Bhattacharyya distance, which appears suboptimal here compared to using a simple Euclidean distance. We hypothesize that the main issue with this approach is that an admixed population covers a large volume in the PCA space, therefore all distances to this population cluster are small because of the covariance component from the Mahalanobis distance. In contrast, here we propose to directly use the global scale of the PC scores, which is invariant from the cluster scattering. This global scale makes it also more robust to infer ancestry with our method as compared to using relative proportions from k=20 nearest neighbors (kNN, Zhang *et al.* (2020)). Indeed, consider e.g. an admixed individual of say 25% European ancestry and 75% African ancestry. The kNN-based method is likely to identify this individual as of African ancestry, while our method will probably be unable to match it, which is a beneficial feature when we are interested in defining genetically homogeneous groups. We also believe our proposed method to be more robust than machine learning methods, because a machine learning method would try e.g. to differentiate between GBR and CEU 1000G populations, which are two very close populations of Northwest Europe (*F*_*ST*_ of 0.0002). In other words, our distance-based method should benefit from the inclusion of any new reference population while it would make it increasingly complex to apply machine learning methods.

Yet, our proposed method also has limitations. First, since we match target individuals to 1000G populations, if individuals are far from all 26 1000G populations, then they would not be matched. When looking at the POPRES data, more individuals from East Europe could not be matched. This is not surprising because there are no East European population in the 1000G data. Moreover, if we look at the location of the 1000G populations on a map, we can see that it lacks representation of many parts of the world (Figure 3). This issue has also been reported e.g. for Asian populations (Lu and Xu 2013). Therefore more diverse populations should be aggregated to better cover the worldwide genome diversity, which would also improve our proposed method. Nevertheless, we also show how to define homogeneous ancestry groups without using the 1000G data, either by using self-reported ancestries or countries of birth. When a predominant genetic ancestry is present in the data, such as British in the UK Biobank (Bycroft *et al.* 2018) or Danish in the iPSYCH data (Pedersen *et al.* 2018), we also show how to directly restrict to a homogeneous subset of the data.

**Figure 3:**
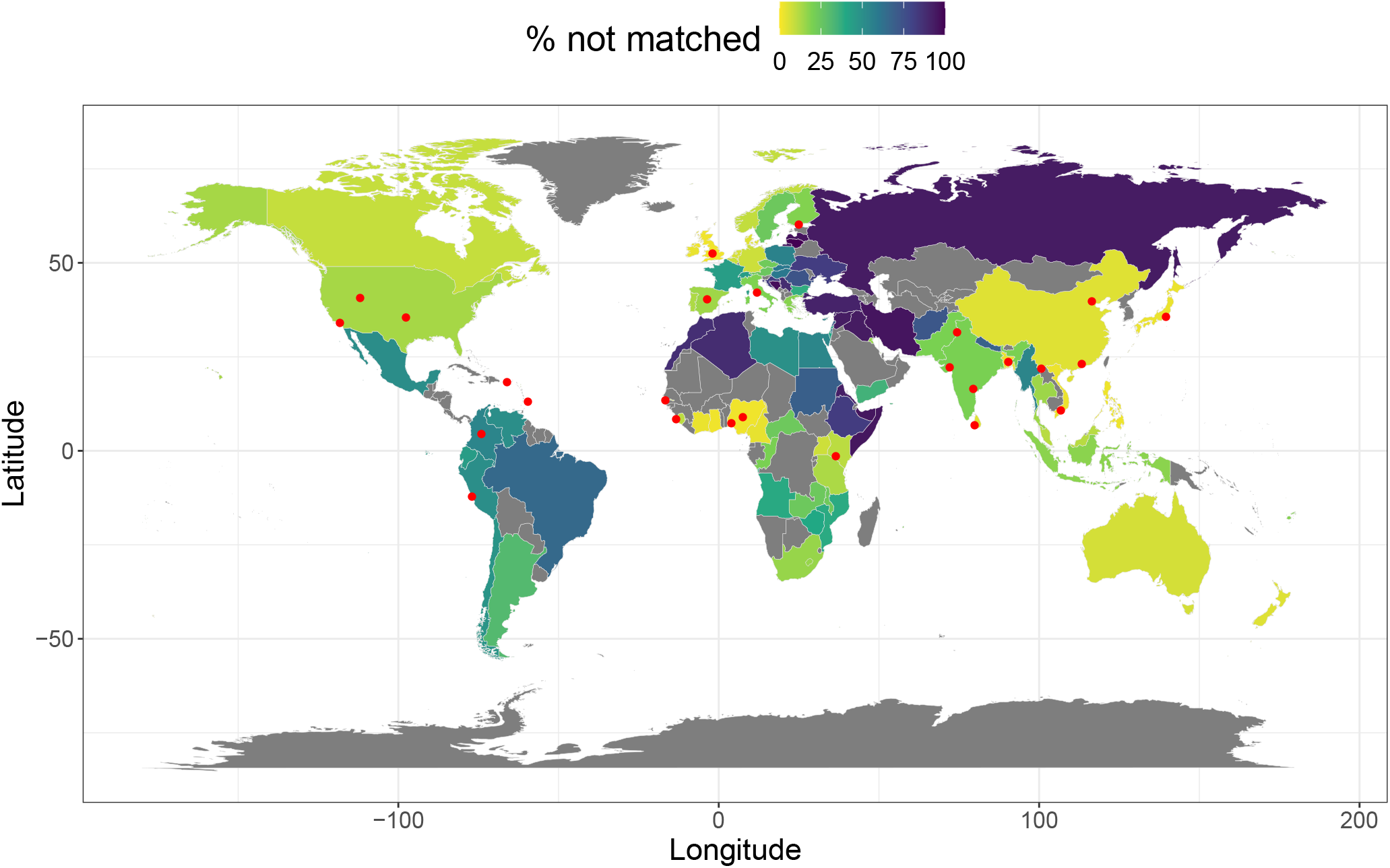
Percentage of individuals from the UK Biobank that could not been matched to any of the 26 1000G populations using our method, per country of birth (Field 20115). Countries in grey contain less than 30 individuals, therefore their percentages are not represented. Red points represent the locations of the 1000G populations, accessed from https://www.internationalgenome.org/data-portal/population. Note that “Gujarati Indian from Houston, Texas” were manually moved to Gujarat (22.309425, 72.136230), “Sri Lankan Tamil from the UK” to Sri Lanka (6.927079, 79.861244), and “Indian Telugu from the UK” to (16.5, 79.5) to better reflect the location of their ancestors. Also note that “Utah Residents with Northern and Western European Ancestry”, “Americans of African Ancestry in SW USA”, “African Caribbeans in Barbados” and “Mexican Ancestry from Los Angeles USA” are probably not located at their ancestral location.

A second potential limitation of our method is that it has two hyper-parameters: the number of PCs used to compute the distances and the threshold on the minimum distance to any cluster center above which the ancestry is not matched. Several studies have used only the first two PCs for ancestry inference. We have shown here that using two PCs (or even four) is not enough for distinguishing between populations at the sub-continental level (Figure S5). As in Privé *et al.* (2020), we recommend to use all PCs that visually separate some populations. Moreover, we believe our proposed method to be robust to increasing the number of PCs used because contribution to the Euclidean distance is smaller for later PCs than for first PCs. As for the distance limit, we have shown here how to define it to approximately correspond to an *F*_*ST*_ of 0.002. Alternatively, a threshold can be chosen based on the visual inspection of the histogram of distances (on a log scale). This threshold can also be adjusted depending on how homogeneous one want each cluster to be.

In conclusion, we believe our proposed approach to be a simple and robust way to infer global ancestry and to define groups of homogeneous ancestry. It is also very fast, allowing to infer ancestry for 488,371 individuals in 20 minutes using 16 cores.

## Software and code availability

The newest version of R package bigsnpr can be installed from GitHub (see https://github.com/privefl/bigsnpr). The code used in this paper is available at https://github.com/privefl/paper-ancestry-matching/tree/master/code.

## Acknowledgements

Authors thank Alex Diaz-Papkovich, Clive Hoggart and others for their useful feedback. This research has been conducted using the UK Biobank Resource under Application Number 41181.

## Funding

F.P. is supported by the Danish National Research Foundation (Niels Bohr Professorship to John McGrath).

## Declaration of Interests

The authors declare no competing interests.

## Supplementary Materials

### Definitions † and methods

- The **1000 Genomes Project (1000G)** data is composed of approximately 100 individuals for each of 26 populations worldwide (described at https://www.internationalgenome.org/category/population/), including 7 African (AFR), 5 East Asian (EAS), 5 South Asian (SAS), 5 European (EUR) and 4 admixed American (AMR) populations. Here we used the transformed data in PLINK format provided in Privé *et al.* (2020).
- The ***F***_***ST***_ measures the relative amount of genetic variance between populations compared to the total genetic variance within these populations (Wright 1965). We use the weighted average formula proposed in Weir and Cockerham (1984), which we now implement in our package bigsnpr (Privé *et al.* 2018).
- The **Principal Component (PC) scores** are defined as *U* Δ, where *U* Δ*V*^*T*^ is the singular value decomposition of the (scaled) genotype matrix (Privé *et al.* 2020). They are usually truncated, e.g. corresponding to the first 20 principal dimensions only.
- The **Bhattacharyya distance** between two multivariate normal distributions 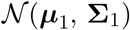 and 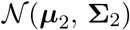 is defined as 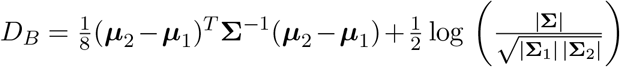, where 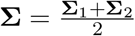 and |*M*| is the absolute value of the determinant of matrix *M* (Bhattacharyya 1943; Fukunaga 1990). The mean and covariance parameters for each population are computed using the robust location and covariance parameters as proposed in Privé *et al.* (2020).
- The **geometric median** of points is the point that minimizes the sum of all Euclidean distances to these points. We now implement this as function `geometric_median` in our R package bigutilsr.
- The **UK Biobank** is a large cohort of half a million individuals from the UK, for which we have access to both genotypes and multiple phenotypes (https://www.ukbiobank.ac.uk/). We apply some quality control filters to the genotyped data; we remove individuals with more than 10% missing values, variants with more than 1% missing values, variants having a minor allele frequency < 0.01, variants with P-value of the Hardy-Weinberg exact test < 10^*−*50^, and non-autosomal variants. This results in 488,371 individuals and 504,139 genetic variants.

### Measures of genetic dissimilarity between populations

**Table S1:** *F*_*ST*_ values between African populations of the 1000G and all 26 1000G populations.

|  | LWK | ESN | YRI | ACB | ASW | GWD | MSL |
| --- | --- | --- | --- | --- | --- | --- | --- |
| LWK | 0.0000 | 0.0077 | 0.0071 | 0.0064 | 0.0090 | 0.0108 | 0.0093 |
| ESN | 0.0077 | 0.0000 | 0.0008 | 0.0034 | 0.0088 | 0.0075 | 0.0051 |
| YRI | 0.0071 | 0.0008 | 0.0000 | 0.0025 | 0.0080 | 0.0062 | 0.0039 |
| ACB | 0.0064 | 0.0034 | 0.0025 | 0.0000 | 0.0020 | 0.0060 | 0.0044 |
| ASW | 0.0090 | 0.0088 | 0.0080 | 0.0020 | 0.0000 | 0.0098 | 0.0094 |
| GWD | 0.0108 | 0.0075 | 0.0062 | 0.0060 | 0.0098 | 0.0000 | 0.0036 |
| MSL | 0.0093 | 0.0051 | 0.0039 | 0.0044 | 0.0094 | 0.0036 | 0.0000 |
| JPT | 0.1475 | 0.1564 | 0.1545 | 0.1344 | 0.1194 | 0.1517 | 0.1574 |
| CHB | 0.1456 | 0.1546 | 0.1527 | 0.1324 | 0.1174 | 0.1499 | 0.1556 |
| CHS | 0.1466 | 0.1555 | 0.1536 | 0.1335 | 0.1186 | 0.1509 | 0.1565 |
| CDX | 0.1456 | 0.1544 | 0.1526 | 0.1324 | 0.1178 | 0.1498 | 0.1555 |
| KHV | 0.1435 | 0.1525 | 0.1507 | 0.1304 | 0.1154 | 0.1479 | 0.1535 |
| GIH | 0.1101 | 0.1200 | 0.1186 | 0.0954 | 0.0773 | 0.1156 | 0.1200 |
| PJL | 0.1069 | 0.1167 | 0.1154 | 0.0920 | 0.0735 | 0.1124 | 0.1167 |
| BEB | 0.1077 | 0.1174 | 0.1161 | 0.0934 | 0.0755 | 0.1131 | 0.1174 |
| ITU | 0.1096 | 0.1195 | 0.1181 | 0.0954 | 0.0778 | 0.1151 | 0.1195 |
| STU | 0.1091 | 0.1189 | 0.1175 | 0.0949 | 0.0774 | 0.1145 | 0.1189 |
| PEL | 0.1472 | 0.1559 | 0.1541 | 0.1325 | 0.1144 | 0.1515 | 0.1567 |
| MXL | 0.1125 | 0.1219 | 0.1205 | 0.0972 | 0.0772 | 0.1175 | 0.1218 |
| CLM | 0.0970 | 0.1063 | 0.1051 | 0.0816 | 0.0620 | 0.1021 | 0.1061 |
| PUR | 0.0849 | 0.0938 | 0.0927 | 0.0699 | 0.0515 | 0.0898 | 0.0935 |
| FIN | 0.1219 | 0.1319 | 0.1306 | 0.1044 | 0.0837 | 0.1272 | 0.1319 |
| CEU | 0.1189 | 0.1291 | 0.1278 | 0.1014 | 0.0805 | 0.1244 | 0.1290 |
| GBR | 0.1193 | 0.1295 | 0.1282 | 0.1017 | 0.0808 | 0.1248 | 0.1294 |
| IBS | 0.1145 | 0.1247 | 0.1234 | 0.0975 | 0.0772 | 0.1199 | 0.1247 |
| TSI | 0.1154 | 0.1258 | 0.1245 | 0.0986 | 0.0783 | 0.1210 | 0.1258 |

**Table S2:** *F*_*ST*_ values between admixed American populations of the 1000G and all 26 1000G populations.

|  | PEL | MXL | CLM | PUR |
| --- | --- | --- | --- | --- |
| LWK | 0.1472 | 0.1125 | 0.0970 | 0.0849 |
| ESN | 0.1559 | 0.1219 | 0.1063 | 0.0938 |
| YRI | 0.1541 | 0.1205 | 0.1051 | 0.0927 |
| ACB | 0.1325 | 0.0972 | 0.0816 | 0.0699 |
| ASW | 0.1144 | 0.0772 | 0.0620 | 0.0515 |
| GWD | 0.1515 | 0.1175 | 0.1021 | 0.0898 |
| MSL | 0.1567 | 0.1218 | 0.1061 | 0.0935 |
| JPT | 0.0795 | 0.0643 | 0.0707 | 0.0773 |
| CHB | 0.0786 | 0.0628 | 0.0689 | 0.0752 |
| CHS | 0.0811 | 0.0650 | 0.0708 | 0.0769 |
| CDX | 0.0849 | 0.0675 | 0.0719 | 0.0773 |
| KHV | 0.0817 | 0.0643 | 0.0689 | 0.0744 |
| GIH | 0.0725 | 0.0370 | 0.0278 | 0.0269 |
| PJL | 0.0688 | 0.0327 | 0.0230 | 0.0220 |
| BEB | 0.0669 | 0.0344 | 0.0278 | 0.0282 |
| ITU | 0.0732 | 0.0391 | 0.0308 | 0.0303 |
| STU | 0.0728 | 0.0390 | 0.0309 | 0.0305 |
| PEL | 0.0000 | 0.0170 | 0.0380 | 0.0548 |
| MXL | 0.0170 | 0.0000 | 0.0090 | 0.0180 |
| CLM | 0.0380 | 0.0090 | 0.0000 | 0.0056 |
| PUR | 0.0548 | 0.0180 | 0.0056 | 0.0000 |
| FIN | 0.0772 | 0.0338 | 0.0178 | 0.0149 |
| CEU | 0.0804 | 0.0334 | 0.0143 | 0.0100 |
| GBR | 0.0809 | 0.0338 | 0.0146 | 0.0102 |
| IBS | 0.0820 | 0.0339 | 0.0134 | 0.0081 |
| TSI | 0.0825 | 0.0345 | 0.0143 | 0.0090 |

**Table S3:** *F*_*ST*_ values between East Asian populations of the 1000G and all 26 1000G populations.

|  | JPT | CHB | CHS | CDX | KHV |
| --- | --- | --- | --- | --- | --- |
| LWK | 0.1475 | 0.1456 | 0.1466 | 0.1456 | 0.1435 |
| ESN | 0.1564 | 0.1546 | 0.1555 | 0.1544 | 0.1525 |
| YRI | 0.1545 | 0.1527 | 0.1536 | 0.1526 | 0.1507 |
| ACB | 0.1344 | 0.1324 | 0.1335 | 0.1324 | 0.1304 |
| ASW | 0.1194 | 0.1174 | 0.1186 | 0.1178 | 0.1154 |
| GWD | 0.1517 | 0.1499 | 0.1509 | 0.1498 | 0.1479 |
| MSL | 0.1574 | 0.1556 | 0.1565 | 0.1555 | 0.1535 |
| JPT | 0.0000 | 0.0068 | 0.0086 | 0.0166 | 0.0140 |
| CHB | 0.0068 | 0.0000 | 0.0010 | 0.0084 | 0.0062 |
| CHS | 0.0086 | 0.0010 | 0.0000 | 0.0047 | 0.0031 |
| CDX | 0.0166 | 0.0084 | 0.0047 | 0.0000 | 0.0016 |
| KHV | 0.0140 | 0.0062 | 0.0031 | 0.0016 | 0.0000 |
| GIH | 0.0693 | 0.0673 | 0.0685 | 0.0685 | 0.0650 |
| PJL | 0.0669 | 0.0647 | 0.0660 | 0.0660 | 0.0626 |
| BEB | 0.0542 | 0.0518 | 0.0528 | 0.0527 | 0.0494 |
| ITU | 0.0656 | 0.0636 | 0.0647 | 0.0646 | 0.0611 |
| STU | 0.0642 | 0.0623 | 0.0634 | 0.0633 | 0.0598 |
| PEL | 0.0795 | 0.0786 | 0.0811 | 0.0849 | 0.0817 |
| MXL | 0.0643 | 0.0628 | 0.0650 | 0.0675 | 0.0643 |
| CLM | 0.0707 | 0.0689 | 0.0708 | 0.0719 | 0.0689 |
| PUR | 0.0773 | 0.0752 | 0.0769 | 0.0773 | 0.0744 |
| FIN | 0.0924 | 0.0901 | 0.0920 | 0.0925 | 0.0893 |
| CEU | 0.0985 | 0.0960 | 0.0977 | 0.0978 | 0.0946 |
| GBR | 0.0993 | 0.0968 | 0.0985 | 0.0985 | 0.0953 |
| IBS | 0.0981 | 0.0957 | 0.0973 | 0.0973 | 0.0942 |
| TSI | 0.0981 | 0.0956 | 0.0972 | 0.0972 | 0.0940 |

**Table S4:** *F*_*ST*_ values between European populations of the 1000G and all 26 1000G populations.

|  | FIN | CEU | GBR | IBS | TSI |
| --- | --- | --- | --- | --- | --- |
| LWK | 0.1219 | 0.1189 | 0.1193 | 0.1145 | 0.1154 |
| ESN | 0.1319 | 0.1291 | 0.1295 | 0.1247 | 0.1258 |
| YRI | 0.1306 | 0.1278 | 0.1282 | 0.1234 | 0.1245 |
| ACB | 0.1044 | 0.1014 | 0.1017 | 0.0975 | 0.0986 |
| ASW | 0.0837 | 0.0805 | 0.0808 | 0.0772 | 0.0783 |
| GWD | 0.1272 | 0.1244 | 0.1248 | 0.1199 | 0.1210 |
| MSL | 0.1319 | 0.1290 | 0.1294 | 0.1247 | 0.1258 |
| JPT | 0.0924 | 0.0985 | 0.0993 | 0.0981 | 0.0981 |
| CHB | 0.0901 | 0.0960 | 0.0968 | 0.0957 | 0.0956 |
| CHS | 0.0920 | 0.0977 | 0.0985 | 0.0973 | 0.0972 |
| CDX | 0.0925 | 0.0978 | 0.0985 | 0.0973 | 0.0972 |
| KHV | 0.0893 | 0.0946 | 0.0953 | 0.0942 | 0.0940 |
| GIH | 0.0343 | 0.0325 | 0.0328 | 0.0334 | 0.0317 |
| PJL | 0.0289 | 0.0269 | 0.0272 | 0.0278 | 0.0262 |
| BEB | 0.0372 | 0.0368 | 0.0372 | 0.0375 | 0.0362 |
| ITU | 0.0393 | 0.0380 | 0.0384 | 0.0384 | 0.0367 |
| STU | 0.0398 | 0.0385 | 0.0389 | 0.0389 | 0.0373 |
| PEL | 0.0772 | 0.0804 | 0.0809 | 0.0820 | 0.0825 |
| MXL | 0.0338 | 0.0334 | 0.0338 | 0.0339 | 0.0345 |
| CLM | 0.0178 | 0.0143 | 0.0146 | 0.0134 | 0.0143 |
| PUR | 0.0149 | 0.0100 | 0.0102 | 0.0081 | 0.0090 |
| FIN | 0.0000 | 0.0062 | 0.0066 | 0.0101 | 0.0116 |
| CEU | 0.0062 | 0.0000 | 0.0002 | 0.0022 | 0.0034 |
| GBR | 0.0066 | 0.0002 | 0.0000 | 0.0024 | 0.0037 |
| IBS | 0.0101 | 0.0022 | 0.0024 | 0.0000 | 0.0015 |
| TSI | 0.0116 | 0.0034 | 0.0037 | 0.0015 | 0.0000 |

**Table S5:** *F*_*ST*_ values between South Asian populations of the 1000G and all 26 1000G populations.

|  | GIH | PJL | BEB | ITU | STU |
| --- | --- | --- | --- | --- | --- |
| LWK | 0.1101 | 0.1069 | 0.1077 | 0.1096 | 0.1091 |
| ESN | 0.1200 | 0.1167 | 0.1174 | 0.1195 | 0.1189 |
| YRI | 0.1186 | 0.1154 | 0.1161 | 0.1181 | 0.1175 |
| ACB | 0.0954 | 0.0920 | 0.0934 | 0.0954 | 0.0949 |
| ASW | 0.0773 | 0.0735 | 0.0755 | 0.0778 | 0.0774 |
| GWD | 0.1156 | 0.1124 | 0.1131 | 0.1151 | 0.1145 |
| MSL | 0.1200 | 0.1167 | 0.1174 | 0.1195 | 0.1189 |
| JPT | 0.0693 | 0.0669 | 0.0542 | 0.0656 | 0.0642 |
| CHB | 0.0673 | 0.0647 | 0.0518 | 0.0636 | 0.0623 |
| CHS | 0.0685 | 0.0660 | 0.0528 | 0.0647 | 0.0634 |
| CDX | 0.0685 | 0.0660 | 0.0527 | 0.0646 | 0.0633 |
| KHV | 0.0650 | 0.0626 | 0.0494 | 0.0611 | 0.0598 |
| GIH | 0.0000 | 0.0035 | 0.0042 | 0.0039 | 0.0043 |
| PJL | 0.0035 | 0.0000 | 0.0035 | 0.0033 | 0.0036 |
| BEB | 0.0042 | 0.0035 | 0.0000 | 0.0022 | 0.0021 |
| ITU | 0.0039 | 0.0033 | 0.0022 | 0.0000 | 0.0013 |
| STU | 0.0043 | 0.0036 | 0.0021 | 0.0013 | 0.0000 |
| PEL | 0.0725 | 0.0688 | 0.0669 | 0.0732 | 0.0728 |
| MXL | 0.0370 | 0.0327 | 0.0344 | 0.0391 | 0.0390 |
| CLM | 0.0278 | 0.0230 | 0.0278 | 0.0308 | 0.0309 |
| PUR | 0.0269 | 0.0220 | 0.0282 | 0.0303 | 0.0305 |
| FIN | 0.0343 | 0.0289 | 0.0372 | 0.0393 | 0.0398 |
| CEU | 0.0325 | 0.0269 | 0.0368 | 0.0380 | 0.0385 |
| GBR | 0.0328 | 0.0272 | 0.0372 | 0.0384 | 0.0389 |
| IBS | 0.0334 | 0.0278 | 0.0375 | 0.0384 | 0.0389 |
| TSI | 0.0317 | 0.0262 | 0.0362 | 0.0367 | 0.0373 |

**Figure S1:**
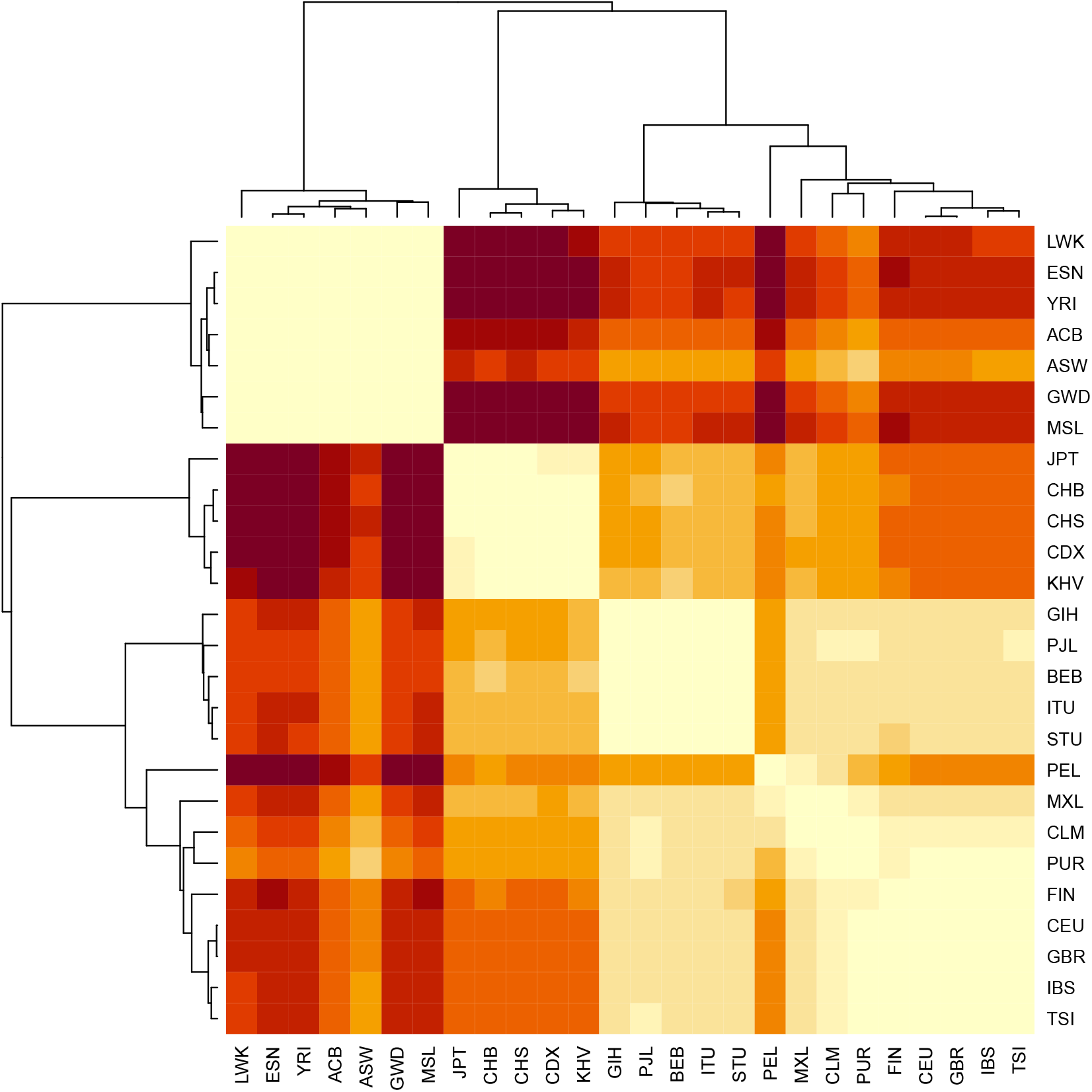
Heatmap with clustering based on the *F*_*ST*_ between pairs of the 26 1000G populations. Corresponding values are reported in tables S1–S5.

**Figure S2:**
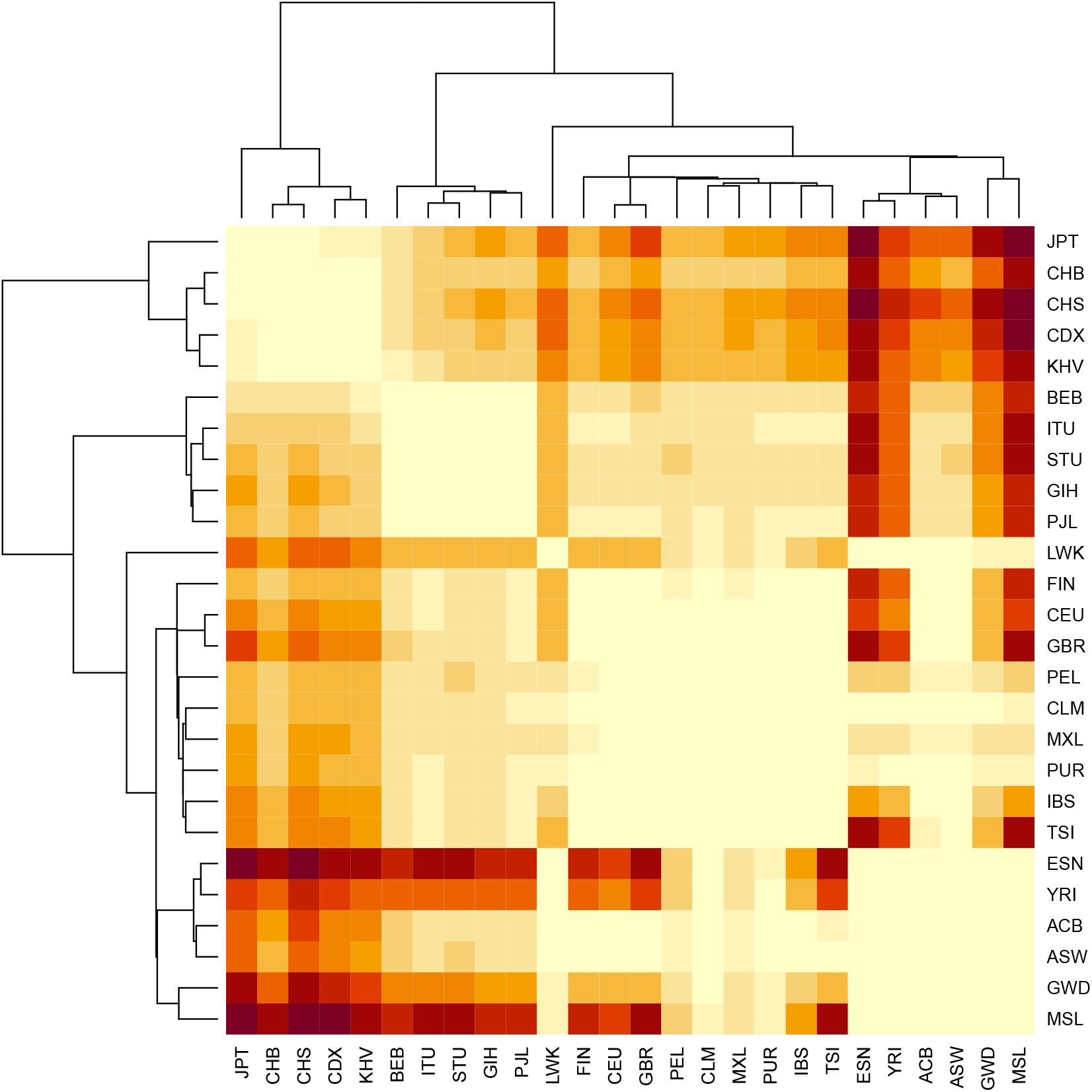
Heatmap with clustering based on the Bhattacharyya distances between pairs of the 26 1000G populations.

**Figure S3:**
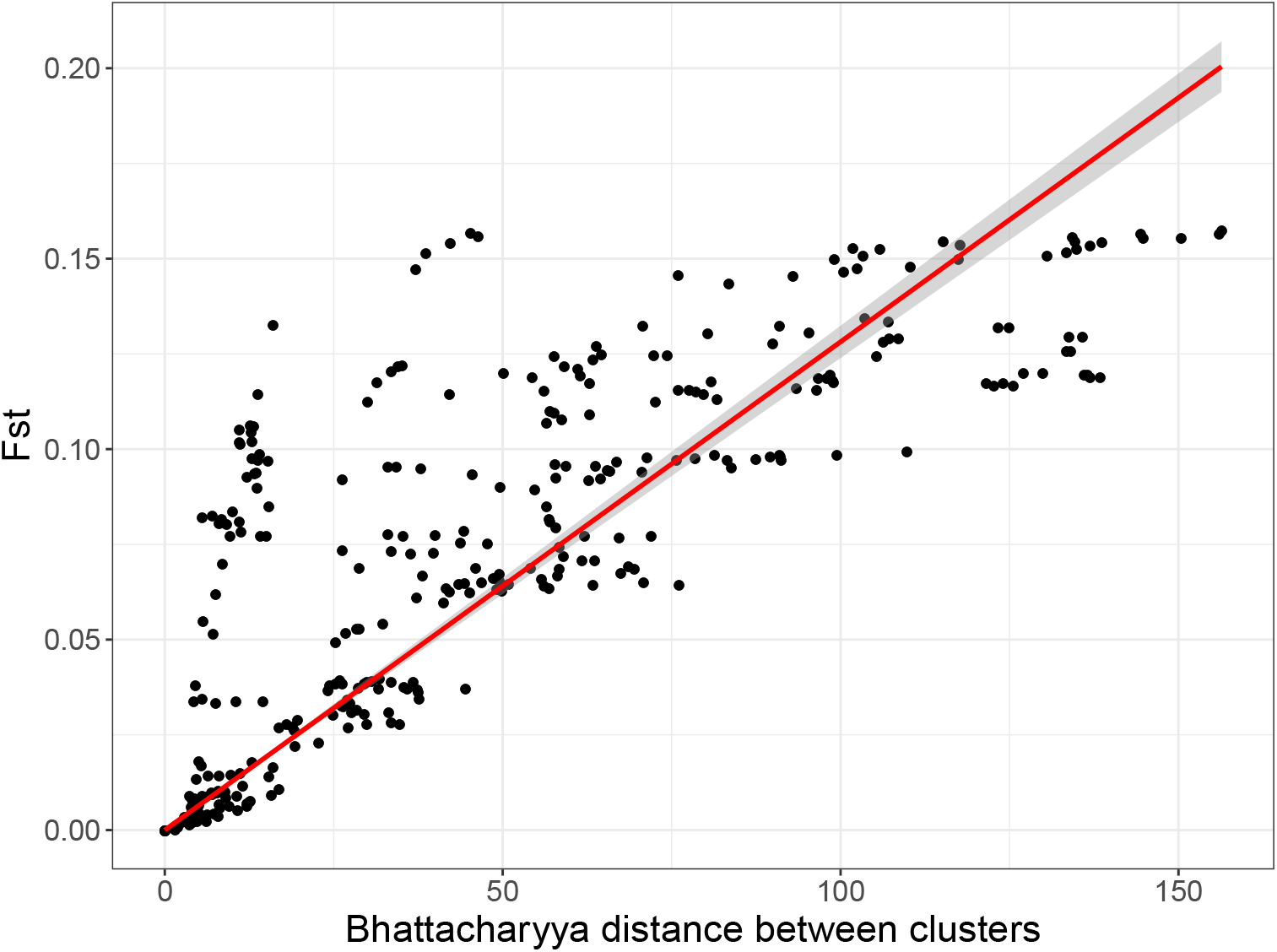
Comparing *F*_*ST*_ to the Bhattacharyya distance on the PCA space between pairs of the 26 1000G populations.

**Figure S4:**
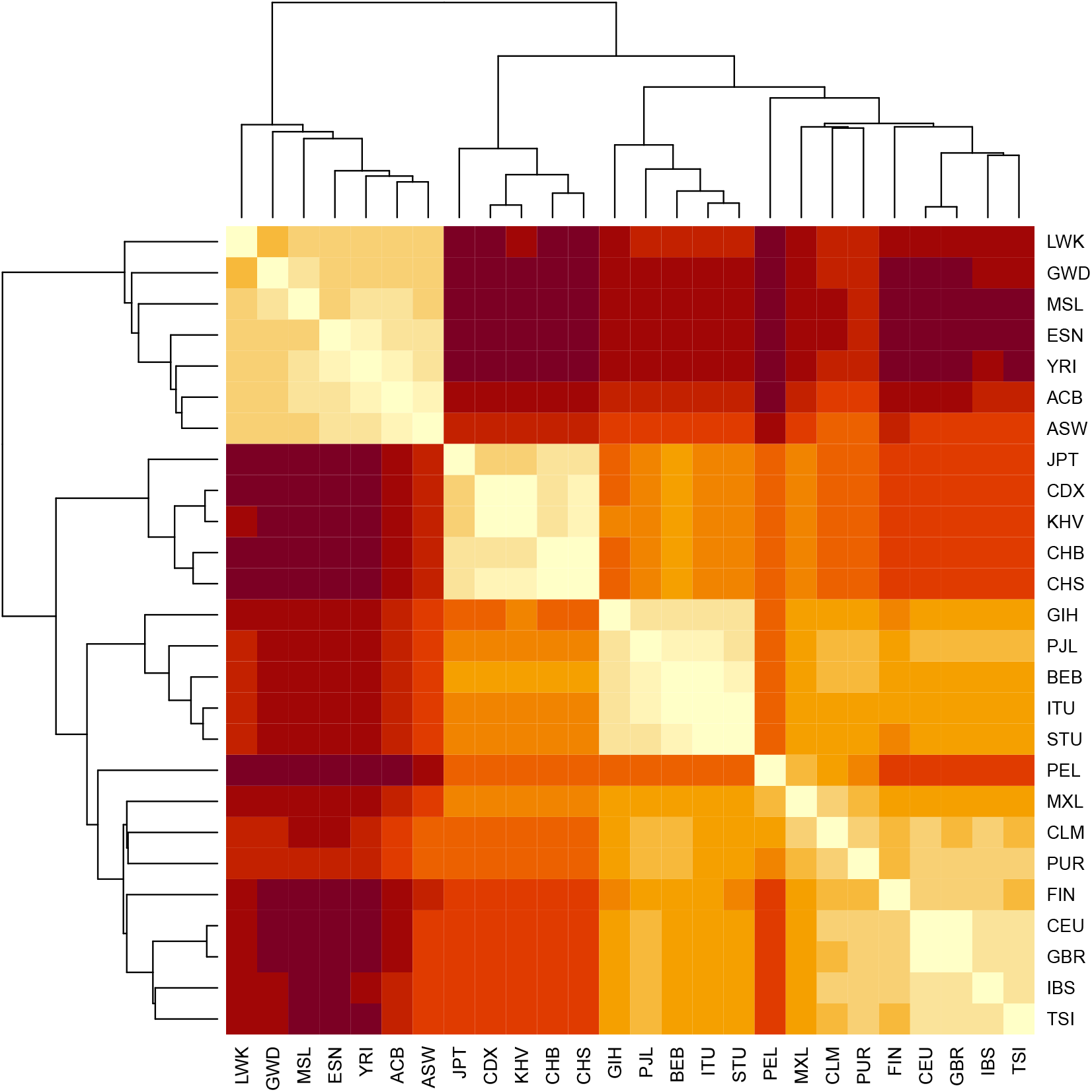
Heatmap with clustering based on the Euclidean distances between centers of pairs of the 26 1000G populations.

**Figure S5:**
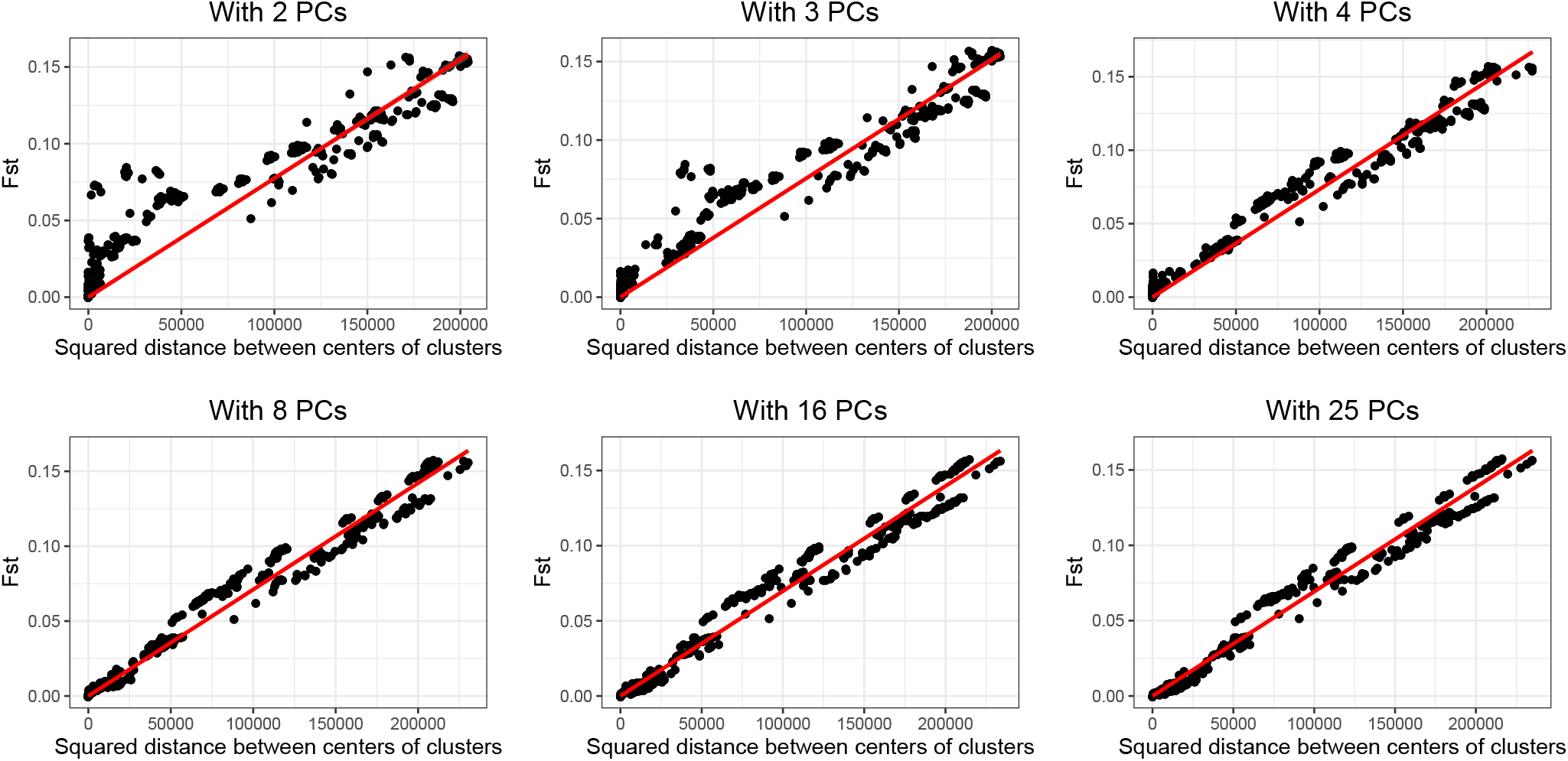
Comparing *F*_*ST*_ to the squared Euclidean distances on the PCA space between centers of pairs of the 26 1000G populations. Distances are computed using different numbers of Principal Components (PCs).

**Figure S6:**
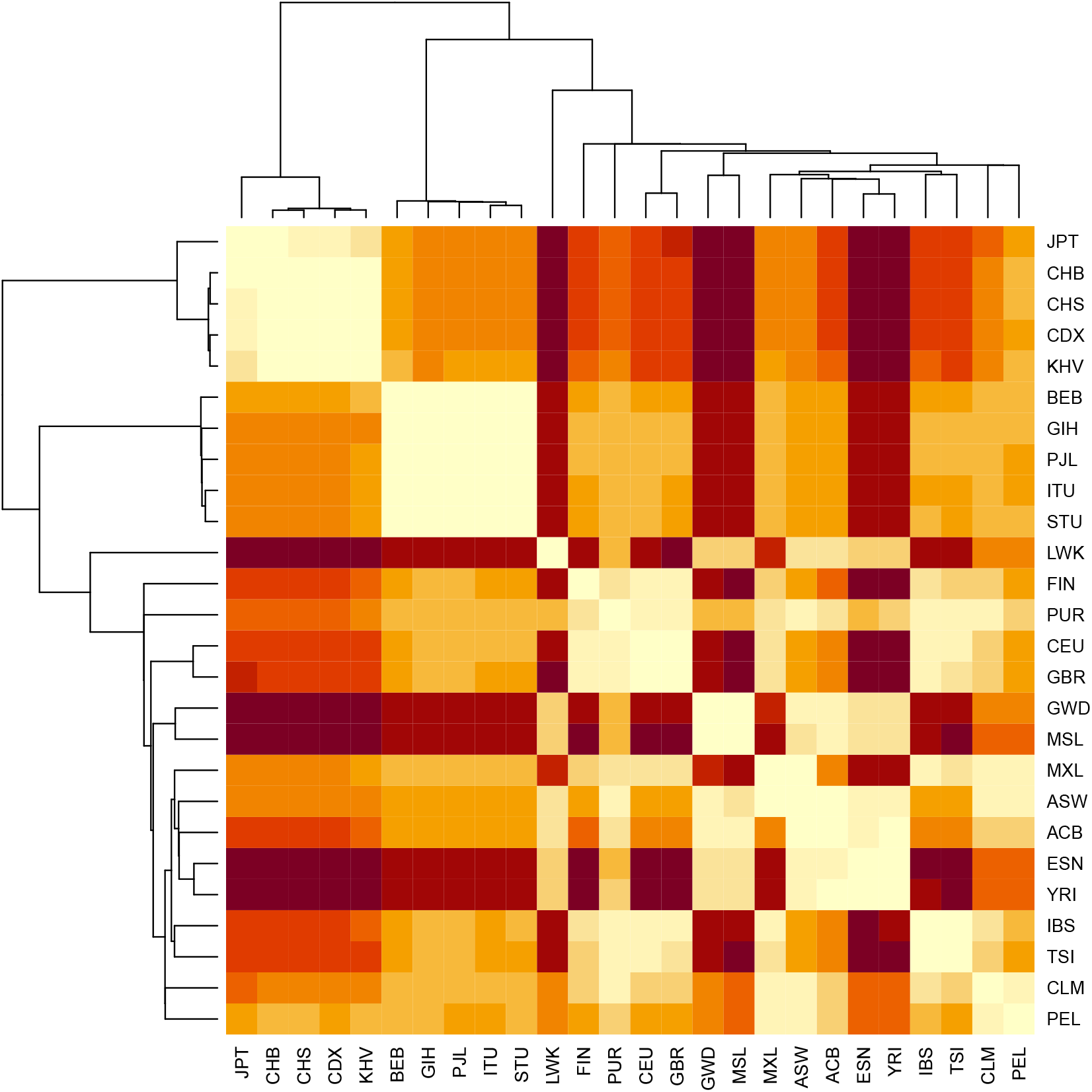
Heatmap with clustering based on the shortest distances between individuals in pairs of the 26 1000G populations.

**Figure S7:**
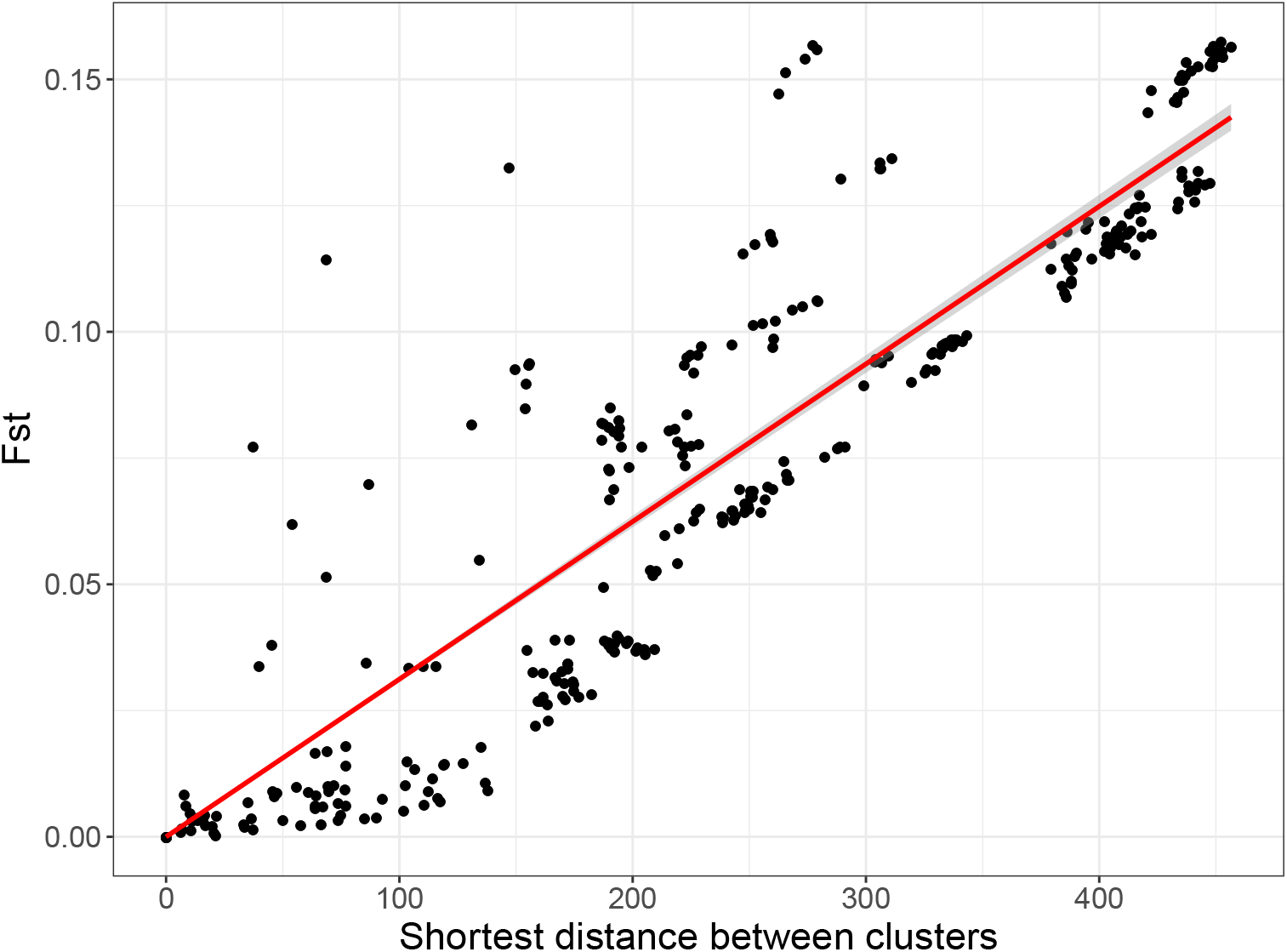
Comparing *F*_*ST*_ to the shortest distances between individuals in pairs of the 26 1000G populations.

### PCA-based ancestry inference

**Table S6:** Self-reported ancestry (left) of UKBB individuals and their matching to 1000G continental populations (top) using 20-wNN. See the description of 1000G populations at https://www.internationalgenome.org/category/population/.

|  | AFR | AMR | EAS | EUR | SAS | Not matched |
| --- | --- | --- | --- | --- | --- | --- |
| British | 4 | 50 | 6 | 430696 | 95 | 163 |
| Irish |  |  |  | 12748 | 3 | 2 |
| White | 1 | 2 | 1 | 540 | 1 |  |
| Other White |  | 170 | 1 | 15533 | 18 | 93 |
| Indian |  |  |  | 21 | 5680 | 15 |
| Pakistani |  |  |  | 3 | 1742 | 3 |
| Bangladeshi |  |  |  |  | 220 | 1 |
| Chinese |  | 7 | 1483 | 3 | 3 | 8 |
| Other Asian | 1 | 1 | 359 | 216 | 1138 | 32 |
| Caribbean | 4117 | 1 |  |  | 36 | 143 |
| African | 3000 |  | 1 | 2 | 2 | 199 |
| Other Black | 90 | 1 |  | 1 | 5 | 21 |
| Asian or Asian British |  |  | 2 | 4 | 34 | 2 |
| Black or Black British | 23 |  |  | 2 |  | 1 |
| White and Black Caribbean | 93 | 16 |  | 74 | 11 | 403 |
| White and Black African | 102 | 13 |  | 52 | 4 | 231 |
| White and Asian |  | 42 | 10 | 242 | 349 | 159 |
| Unknown | 1024 | 541 | 712 | 3774 | 1020 | 749 |

**Table S7:**
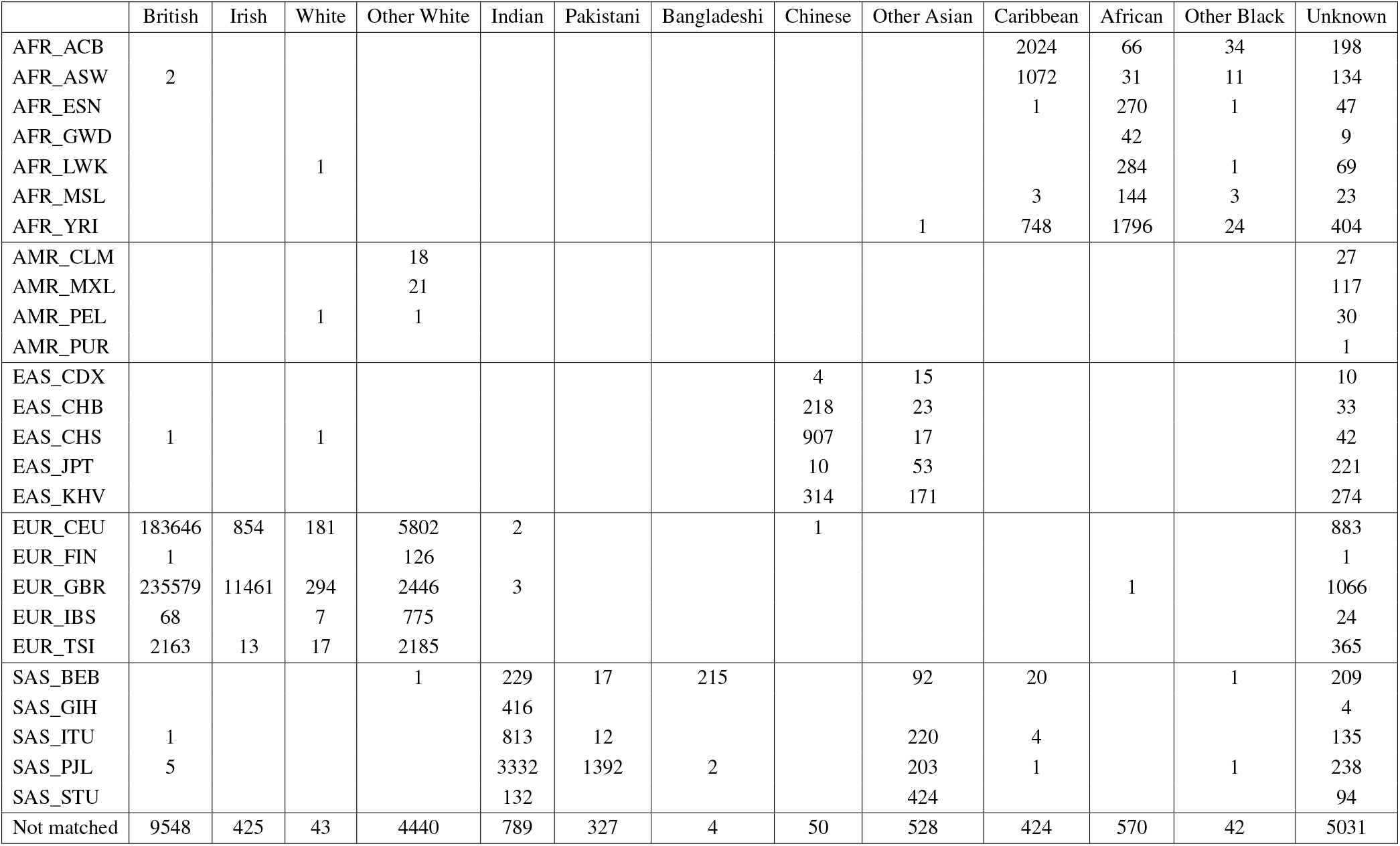
Self-reported ancestry (top) of UKBB individuals and their matching to 1000G populations (left) by our method. See the description of 1000G populations at https://www.internationalgenome.org/category/population/.

|  | British | Irish | White | Other White | Indian | Pakistani | Bangladeshi | Chinese | Other Asian | Caribbean | African | Other Black | Unknown |
| --- | --- | --- | --- | --- | --- | --- | --- | --- | --- | --- | --- | --- | --- |
| AFR_ACB |  |  |  |  |  |  |  |  |  | 2024 | 66 | 34 | 198 |
| AFR_ASW | 2 |  |  |  |  |  |  |  |  | 1072 | 31 | 11 | 134 |
| AFR_ESN |  |  |  |  |  |  |  |  |  | 1 | 270 | 1 | 47 |
| AFR_GWD |  |  |  |  |  |  |  |  |  |  | 42 |  | 9 |
| AFR_LWK |  |  | 1 |  |  |  |  |  |  |  | 284 | 1 | 69 |
| AFR_MSL |  |  |  |  |  |  |  |  |  | 3 | 144 | 3 | 23 |
| AFR_YRI |  |  |  |  |  |  |  |  | 1 | 748 | 1796 | 24 | 404 |
| AMR_CLM |  |  |  | 18 |  |  |  |  |  |  |  |  | 27 |
| AMR_MXL |  |  |  | 21 |  |  |  |  |  |  |  |  | 117 |
| AMR_PEL |  |  | 1 | 1 |  |  |  |  |  |  |  |  | 30 |
| AMR_PUR |  |  |  |  |  |  |  |  |  |  |  |  | 1 |
| EAS_CDX |  |  |  |  |  |  |  | 4 | 15 |  |  |  | 10 |
| EAS_CHB |  |  |  |  |  |  |  | 218 | 23 |  |  |  | 33 |
| EAS_CHS | 1 |  | 1 |  |  |  |  | 907 | 17 |  |  |  | 42 |
| EAS_JPT |  |  |  |  |  |  |  | 10 | 53 |  |  |  | 221 |
| EAS_KHV |  |  |  |  |  |  |  | 314 | 171 |  |  |  | 274 |
| EUR_CEU | 183646 | 854 | 181 | 5802 | 2 |  |  | 1 |  |  |  |  | 883 |
| EUR_FIN | 1 |  |  | 126 |  |  |  |  |  |  |  |  | 1 |
| EUR_GBR | 235579 | 11461 | 294 | 2446 | 3 |  |  |  |  |  | 1 |  | 1066 |
| EUR_IBS | 68 |  | 7 | 775 |  |  |  |  |  |  |  |  | 24 |
| EUR_TSI | 2163 | 13 | 17 | 2185 |  |  |  |  |  |  |  |  | 365 |
| SAS_BEB |  |  |  | 1 | 229 | 17 | 215 |  | 92 | 20 |  | 1 | 209 |
| SAS_GIH |  |  |  |  | 416 |  |  |  |  |  |  |  | 4 |
| SAS_ITU | 1 |  |  |  | 813 | 12 |  |  | 220 | 4 |  |  | 135 |
| SAS_PJL | 5 |  |  |  | 3332 | 1392 | 2 |  | 203 | 1 |  | 1 | 238 |
| SAS_STU |  |  |  |  | 132 |  |  |  | 424 |  |  |  | 94 |
| Not matched | 9548 | 425 | 43 | 4440 | 789 | 327 | 4 | 50 | 528 | 424 | 570 | 42 | 5031 |

**Table S8:** Ancestry (left) of POPRES individuals and their matching to 1000G populations (top) by our method. See the description of 1000G populations at https://www.internationalgenome.org/category/population/.

|  | EUR_CEU | EUR_FIN | EUR_GBR | EUR_IBS | EUR_TSI | Not matched |
| --- | --- | --- | --- | --- | --- | --- |
| Anglo-Irish Isles | 136 |  | 127 |  | 2 | 1 |
| Belgium | 43 |  |  |  |  |  |
| Central Europe | 47 |  |  |  | 8 |  |
| Eastern Europe | 27 |  |  |  | 1 | 2 |
| France | 49 |  | 3 | 35 | 2 |  |
| Germany | 67 |  | 3 |  | 1 |  |
| Italy | 1 |  |  | 11 | 204 | 3 |
| Netherlands | 13 |  | 4 |  |  |  |
| Scandinavia | 13 | 1 | 1 |  |  |  |
| SE Europe | 12 |  |  | 3 | 70 | 9 |
| SW Europe | 1 |  |  | 261 | 1 | 1 |
| Switzerland | 179 |  |  | 32 | 11 |  |

**Figure S8:**
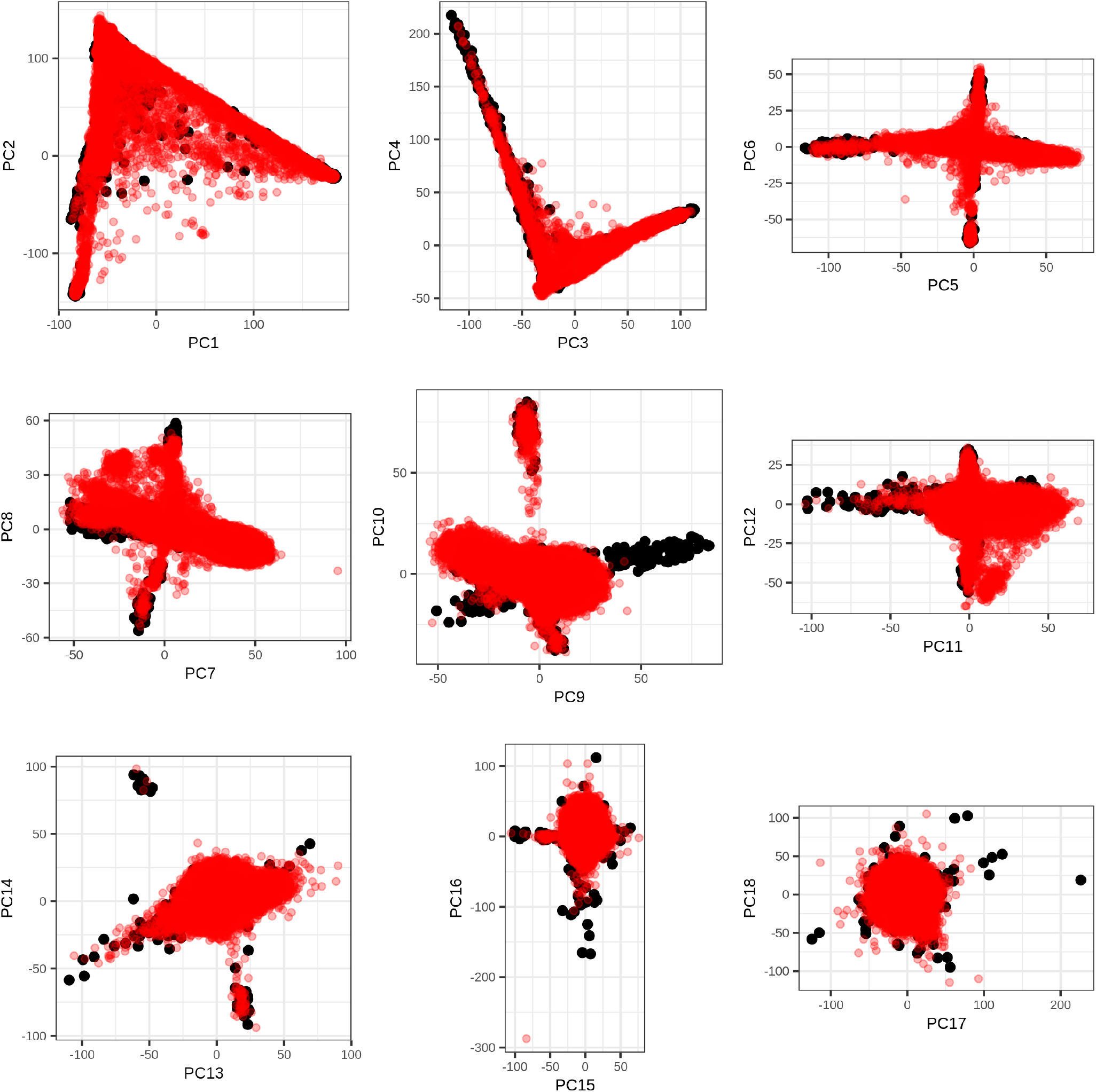
First 18 PC scores of the 1000G data (in black), onto which the UK Biobank data has been projected (in red).

**Figure S9:**
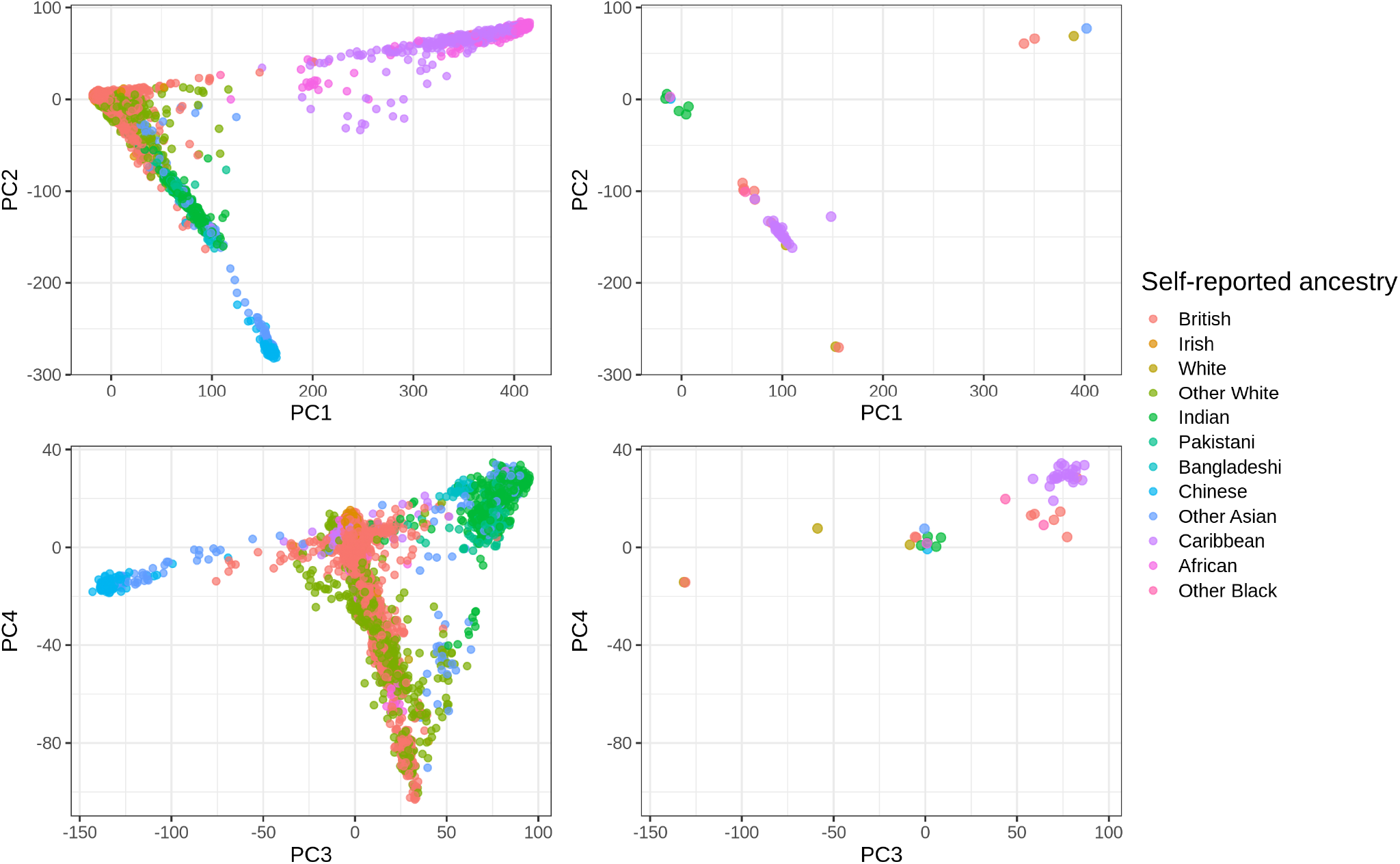
PC scores (computed in the UK Biobank) colored by self-reported ancestry. On the left, these are 50,000 random individuals. On the right, these are the 47 individuals with some discrepancy between their self-reported-ancestry and our ancestry estimation.

### PCA-based ancestry grouping

**Table S9:** Self-reported ancestry (left) of UKBB individuals and their matching to ancestry groups (top) by our method.

|  | British | Indian | Pakistani | Bangladeshi | Chinese | Caribbean | African | Not matched |
| --- | --- | --- | --- | --- | --- | --- | --- | --- |
| British | 426210 | 6 | 4 | 1 | 1 | 2 |  | 4790 |
| Irish | 12712 |  |  |  |  |  |  | 41 |
| White | 492 |  |  |  | 1 |  |  | 52 |
| Other White | 10932 | 1 | 1 | 1 |  |  |  | 4880 |
| Indian | 6 | 1764 | 2488 | 1321 |  |  |  | 137 |
| Pakistani | 1 | 362 | 1299 | 63 |  |  |  | 23 |
| Bangladeshi |  | 3 |  | 215 |  |  |  | 3 |
| Chinese | 1 |  | 1 |  | 1437 |  |  | 65 |
| Other Asian | 4 | 113 | 169 | 745 | 62 |  | 1 | 653 |
| Caribbean |  | 2 |  | 23 |  | 2325 | 1148 | 799 |
| African | 1 |  | 1 |  |  | 74 | 2271 | 857 |
| Other Black |  | 1 | 1 | 1 |  | 36 | 33 | 46 |
| Asian or Asian British |  | 7 | 16 | 3 | 1 |  |  | 15 |
| Black or Black British | 2 |  |  |  |  | 11 | 9 | 4 |
| White and Black Caribbean | 7 |  |  | 1 |  | 10 | 1 | 578 |
| White and Black African | 6 |  |  |  |  | 1 | 2 | 393 |
| White and Asian | 59 | 31 | 7 | 19 |  |  |  | 686 |
| Unknown | 2008 | 129 | 189 | 421 | 114 | 214 | 505 | 4240 |

**Figure S10:**
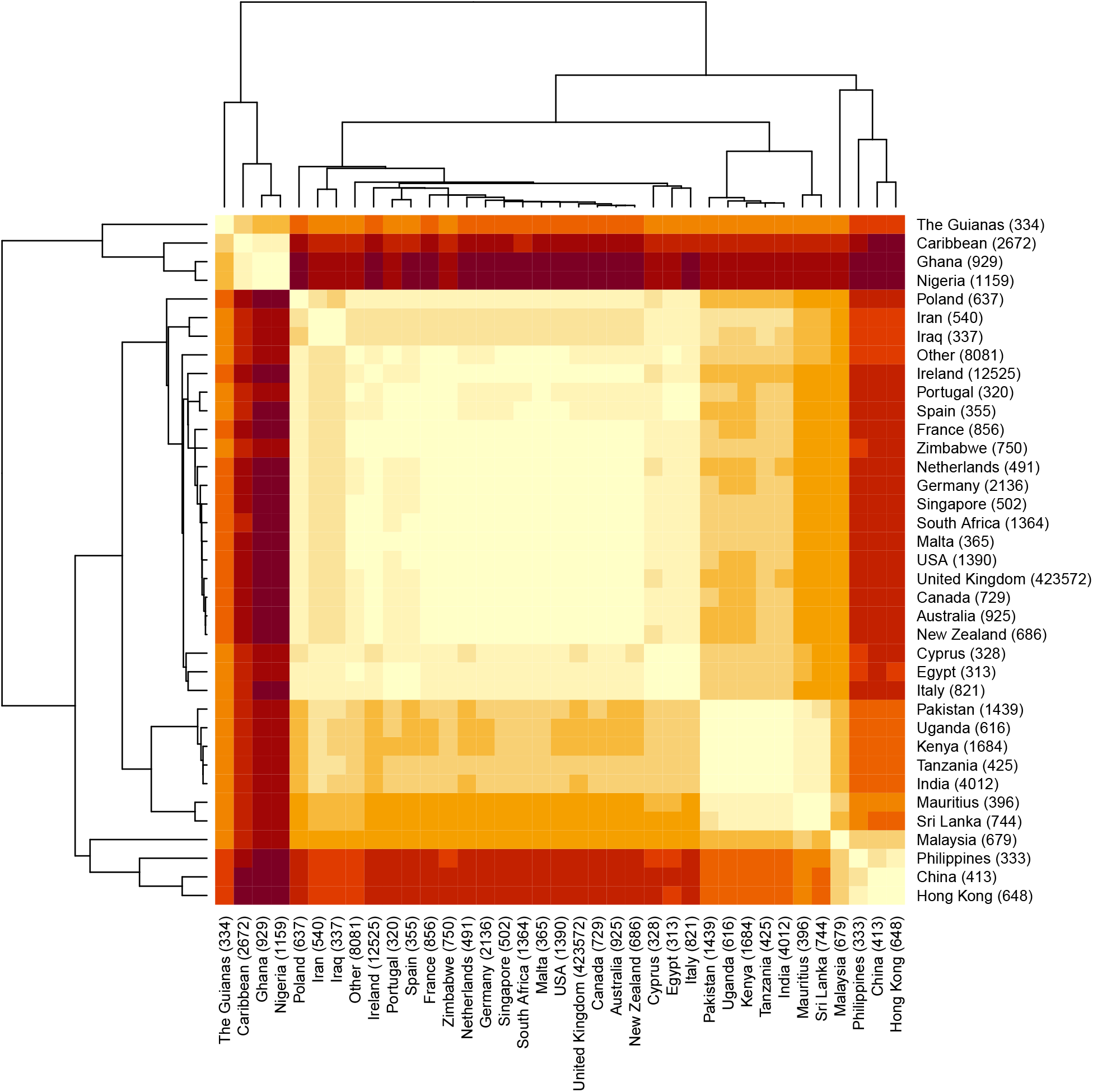
Heatmap with clustering based on the distances in the PCA space between centers of pairs of the countries of birth in the UK Biobank.

**Table S10:**
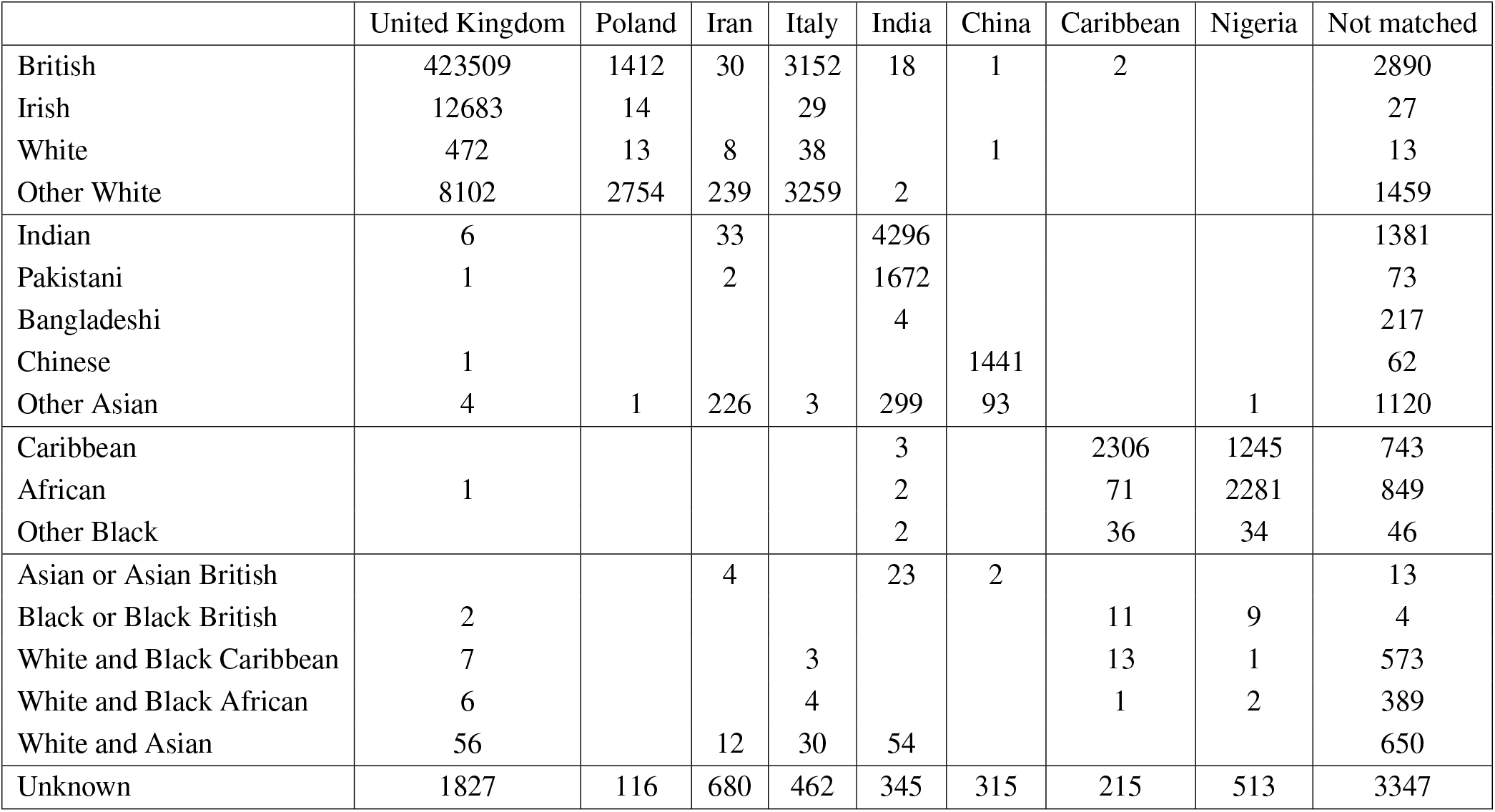
Self-reported ancestry (left) of UKBB individuals and their matching to country groups (top) by our method.

|  | United Kingdom | Poland | Iran | Italy | India | China | Caribbean | Nigeria | Not matched |
| --- | --- | --- | --- | --- | --- | --- | --- | --- | --- |
| British | 423509 | 1412 | 30 | 3152 | 18 | 1 | 2 |  | 2890 |
| Irish | 12683 | 14 |  | 29 |  |  |  |  | 27 |
| White | 472 | 13 | 8 | 38 |  | 1 |  |  | 13 |
| Other White | 8102 | 2754 | 239 | 3259 | 2 |  |  |  | 1459 |
| Indian | 6 |  | 33 |  | 4296 |  |  |  | 1381 |
| Pakistani | 1 |  | 2 |  | 1672 |  |  |  | 73 |
| Bangladeshi |  |  |  |  | 4 |  |  |  | 217 |
| Chinese | 1 |  |  |  |  | 1441 |  |  | 62 |
| Other Asian | 4 | 1 | 226 | 3 | 299 | 93 |  | 1 | 1120 |
| Caribbean |  |  |  |  | 3 |  | 2306 | 1245 | 743 |
| African | 1 |  |  |  | 2 |  | 71 | 2281 | 849 |
| Other Black |  |  |  |  | 2 |  | 36 | 34 | 46 |
| Asian or Asian British |  |  | 4 |  | 23 | 2 |  |  | 13 |
| Black or Black British | 2 |  |  |  |  |  | 11 | 9 | 4 |
| White and Black Caribbean | 7 |  |  | 3 |  |  | 13 | 1 | 573 |
| White and Black African | 6 |  |  | 4 |  |  | 1 | 2 | 389 |
| White and Asian | 56 |  | 12 | 30 | 54 |  |  |  | 650 |
| Unknown | 1827 | 116 | 680 | 462 | 345 | 315 | 215 | 513 | 3347 |

**Figure S11:**
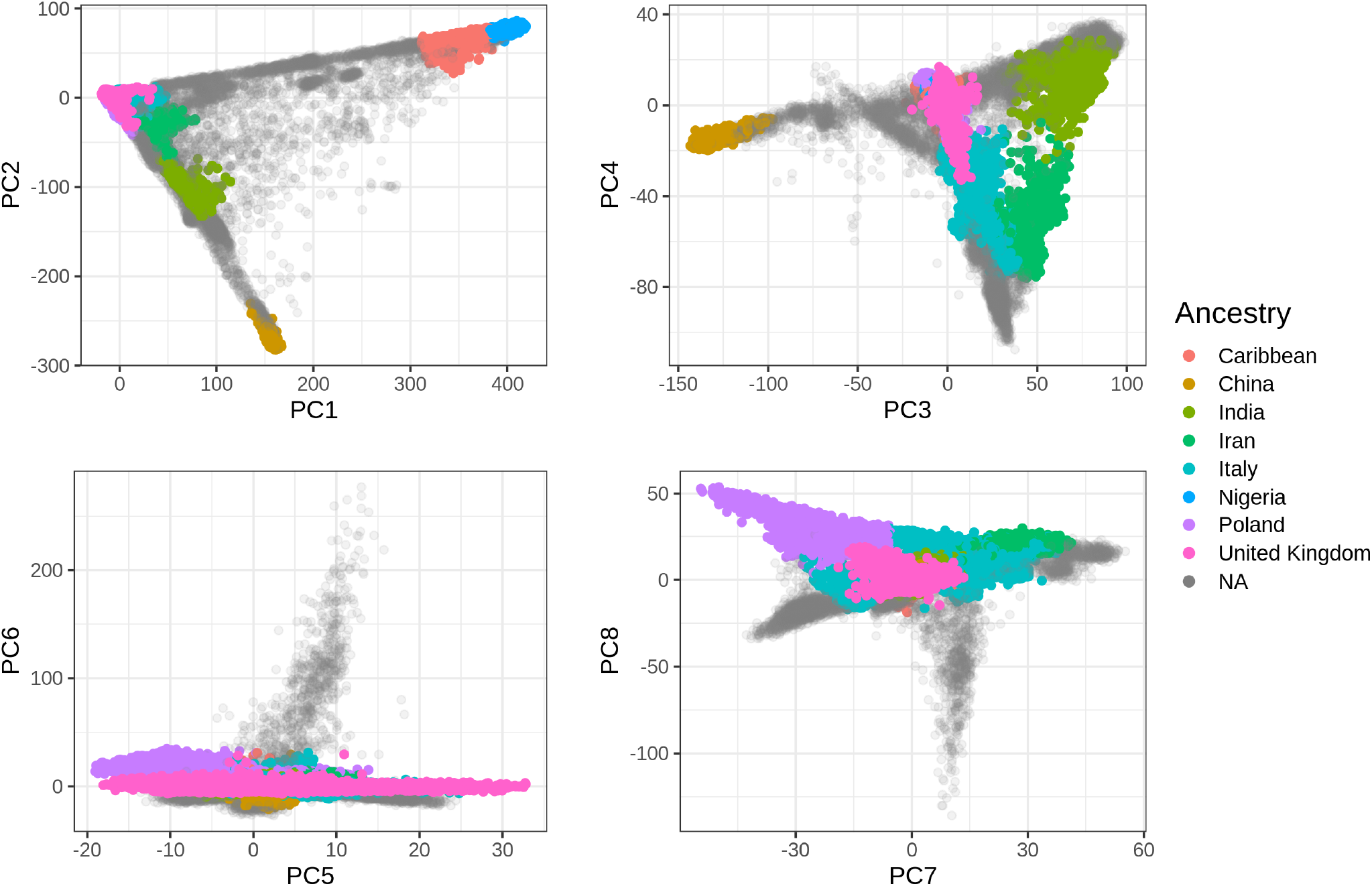
The first eight PC scores computed from the UK Biobank (Field 22009) colored by the homogeneous ancestry group we infer for these individuals.

**Figure S12:**
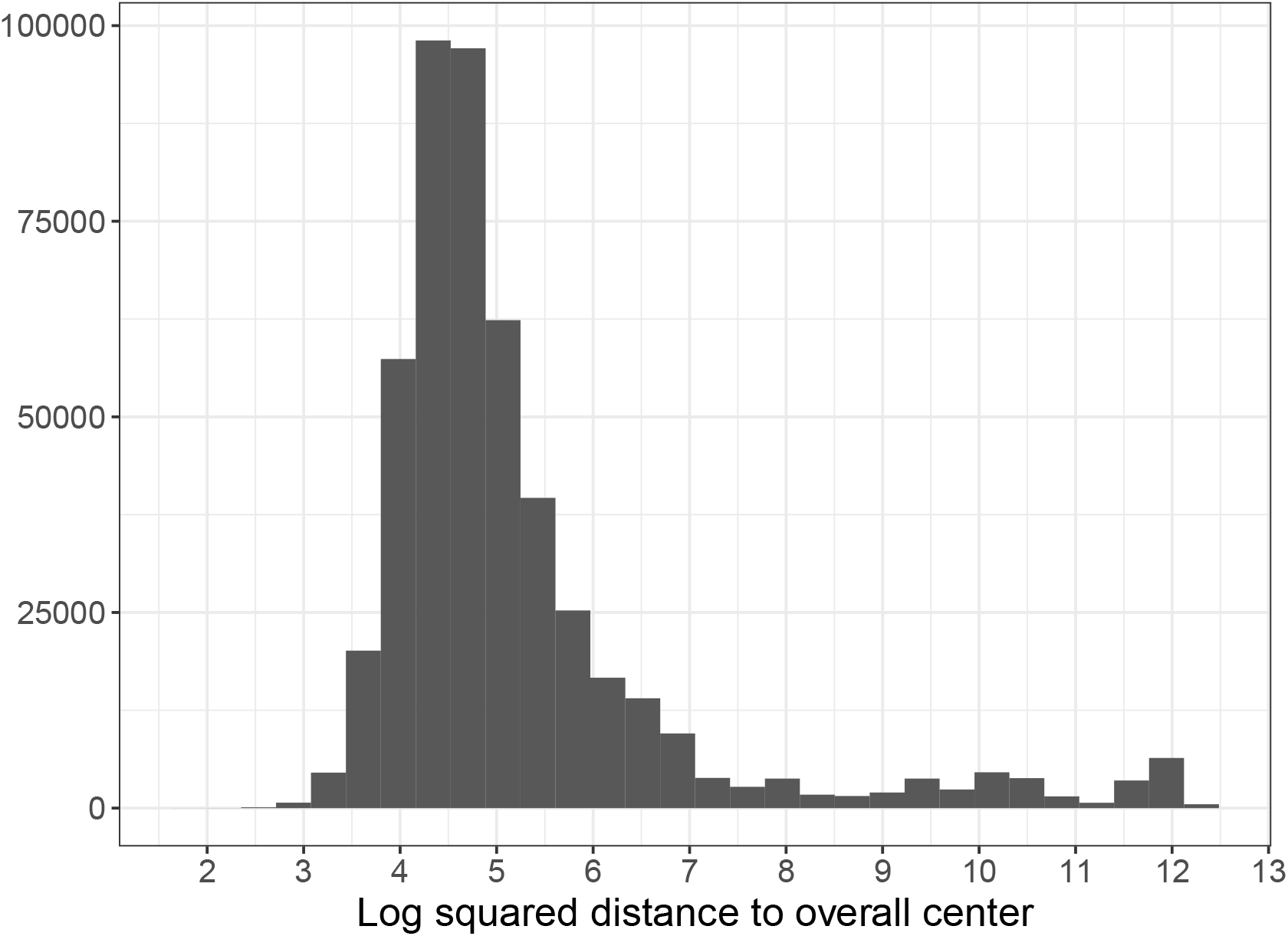
Histogram of (log) squared distances from the UK Biobank PC scores to the geometric median of the all UKBB individuals. Here we use a threshold at 7, based on visual inspection. Alternatively, a more stringent threshold at 6 could also be used.

## Footnotes

† Further defined in supplementary section “Defintions”.

## Notes

### Competing Interest Statement

The authors have declared no competing interest.

### Summary of Updates

- Added some analyses using the countries of birth (Field 20115). - Added a few more analyses, e.g. varying the number of PCs used.

